# Quantitative classification of chromatin dynamics reveals regulators of intestinal stem cell differentiation

**DOI:** 10.1101/637181

**Authors:** Jesse R Raab, Deepthi Y Tulasi, Kortney E Wager, Jeremy M Morowitz, Scott T Magness, Adam D Gracz

## Abstract

Intestinal stem cell (ISC) plasticity is thought to be regulated by broadly-permissive chromatin shared between ISCs and their progeny. Here, we utilize a Sox9^EGFP^ reporter to examine chromatin across ISC differentiation. We find that open chromatin regions (OCRs) can be defined as broadly-permissive or dynamic in a locus-specific manner, with dynamic OCRs found primarily in loci consistent with distal enhancers. By integrating gene expression with chromatin accessibility at transcription factor (TF) motifs in context of Sox9^EGFP^ populations, we classify broadly-permissive and dynamic chromatin relative to TF usage. These analyses identify known and potential regulators of ISC differentiation via their association with dynamic changes in chromatin. We observe ISC expansion in Id3-null mice, consistent with computational predictions. Finally, we examine the relationship between gene expression and 5-hydroxymethylcytosine (5hmC) in Sox9^EGFP^ populations, which reveals 5hmC enrichment in absorptive lineage specific genes. Our data demonstrate that intestinal chromatin dynamics can be quantitatively defined in a locus-specific manner, identify novel potential regulators of ISC differentiation, and provide a chromatin roadmap for further dissecting the role of *cis* regulation of cell fate in the intestine.

## INTRODUCTION

ISCs maintain the intestinal epithelium, which is replaced once a week throughout adult life. To preserve digestive and barrier function, ISCs balance proliferation with differentiation into post-mitotic intestinal epithelial cells (IECs): absorptive enterocytes, goblet cells, Paneth cells, enteroendocrine cells (EECs), and tuft cells. Models for cellular hierarchy in the intestinal epithelium have been complicated by observations of extensive cellular plasticity, wherein secretory and absorptive progenitors as well as post-mitotic Paneth cells can de-differentiate following damage to or ablation of the ISC pool (Buczacki et al., 2013; Schmitt et al., 2018; Tetteh et al., 2016). The genetic mechanisms that allow IECs to adopt metastable differentiated fates while exhibiting facultative ISC function are not fully understood.

Chromatin landscapes consist of histone post-translational modifications, DNA modifications, and higher-order structural organization, and are known to exert regulatory control on cell fate. Chromatin is progressively remodeled during differentiation, and is highly dynamic in embryonic stem cells as well as tissue-specific stem cells in the hair follicle and hematopoietic system (Dixon et al., 2015; Lara-Astiaso et al., 2014; Lien et al., 2011). Recent studies of chromatin in ISCs and differentiated IECs suggest that intestinal chromatin is largely homogenous, with similar enhancer, open chromatin, and DNA methylation profiles observed in ISCs and differentiated progeny (Kaaij et al., 2013; Kim et al., 2014). The observed broadly-permissive nature of chromatin has been proposed as a mechanism for cell fate plasticity in the intestine (Kim et al., 2014). However, these studies relied on pharmacologic or genetic modulation of intestinal epithelial populations to force over-production of absorptive or secretory cells, or have compared *Lgr5+* ISCs to bulk differentiated populations (Barker et al., 2007; Kaaij et al., 2013; Kim et al., 2014). The nature of IEC chromatin has been further complicated by other studies demonstrating histone methylation dynamics between ISCs and differentiated progeny, as well as chromatin remodeling during de-differentiation of facultative ISCs following radiation injury (Jadhav et al., 2017; Kazakevych et al., 2017). The extent to which chromatin is dynamic across ISCs, intermediate transit-amplifying (TA) progenitors, and post-mitotic IECs remains controversial.

Differential expression levels of *Sox9*, observed via a Sox9^EGFP^ BAC transgene, are associated with phenotypically distinct IEC populations (Formeister et al., 2009). Histological and gene expression studies have shown that *Sox9*^low^ cells are consistent with ISCs, *Sox9*^sublow^ cells with TA progenitors, *Sox9*^high^ cells with label-retaining secretory progenitors/facultative ISCs in the crypts and mature EECs and tuft cells in the villus, while *Sox9*^neg^ cells consist of the remaining post-mitotic populations (goblet, Paneth, and absorptive enterocyte) (Formeister et al., 2009; Gracz et al., 2010; Gracz et al., 2015; Roche et al., 2015). We reasoned that this model could be applied to omics-scale assays to understand dynamics in native IEC chromatin. Here, we profile the transcriptome, open chromatin (Omni ATAC-seq), and 5hmC (Nano-hmC-Seal) to interrogate the relationship between gene expression changes and intestinal chromatin dynamics in the Sox9^EGFP^ model. Through the application of quantitative approaches, our analyses demonstrate clear examples of both broadly-permissive and dynamic chromatin in IECs that can be associated with specific transcriptional regulatory networks.

## RESULTS

### *Sox9* expression levels define transcriptomically distinct populations

We isolated *Sox9*^neg^, *Sox9*^sub^, *Sox9*^low^, and *Sox9*^high^ IEC populations from adult Sox9^EGFP^ mice by fluorescence-activated cell sorting (FACS) and subjected them to RNA-seq (Fig. 1A, Fig. S1A), which confirmed that Sox9^EGFP^ levels faithfully recapitulate endogenous *Sox9* expression (Fig. S1B) (Formeister et al., 2009). Previous studies have defined Sox9^EGFP^ populations by RT-qPCR and immunofluorescence, and we next sought to confirm our model with transcriptomic data (Formeister et al., 2009; Gracz et al., 2010). To identify genes enriched in each population, we conducted differential expression analysis and filtered for genes significantly upregulated in a single Sox9^EGFP^ population. We identified 2,852, 253, 1,499, and 3,521 genes that were upregulated in *Sox9*^neg^, *Sox9*^sublow^, *Sox9*^low^, and *Sox9*^high^ populations, respectively (Fig. 1B and Supplemental Table 1). These included genes known to be associated with expected cell types in each Sox9^EGFP^ population: absorptive enterocyte *Fabp1* and goblet cell *Muc5b/6* in *Sox9*^neg^, secretory progenitor *Dll1* in *Sox9*^sublow^, ISC *Ascl2*, *Olfm4*, and *Smoc2* in *Sox9*^low^, EEC *Reg4* and tuft cell *Il25* in *Sox9*^high^ (Fig. 1B). Our data show enrichment of Paneth cell specific *Lyz1/2* in *Sox9*^neg^, consistent with previous observations that the EGFP transgene is silenced in Paneth cells, which express endogenous *Sox9* (Formeister et al., 2009; Gracz et al., 2015). Since our analysis was designed for stringency, Sox9^EGFP^ population gene lists excluded genes expressed highly in more than one population, such as *Lgr5*, which is expressed in *Sox9*^low^ active ISCs as well as *Sox9*^high^ secretory progenitors (Fig. S1C) (Buczacki et al., 2013; Roche et al., 2015). Gene Ontology (GO) analysis identified expected relationships between Sox9^EGFP^ populations and cellular function, including mitosis in *Sox9*^sub^ progenitors, stem cell maintenance in *Sox9*^low^ ISCs, secretory and hormone processes in EEC-enriched *Sox9*^high^, and absorption and secretion in *Sox9*^neg^, which contains absorptive enterocytes as well as secretory goblet and Paneth cells (Fig. S1E and Supplemental Table 2).

**Figure 1:**
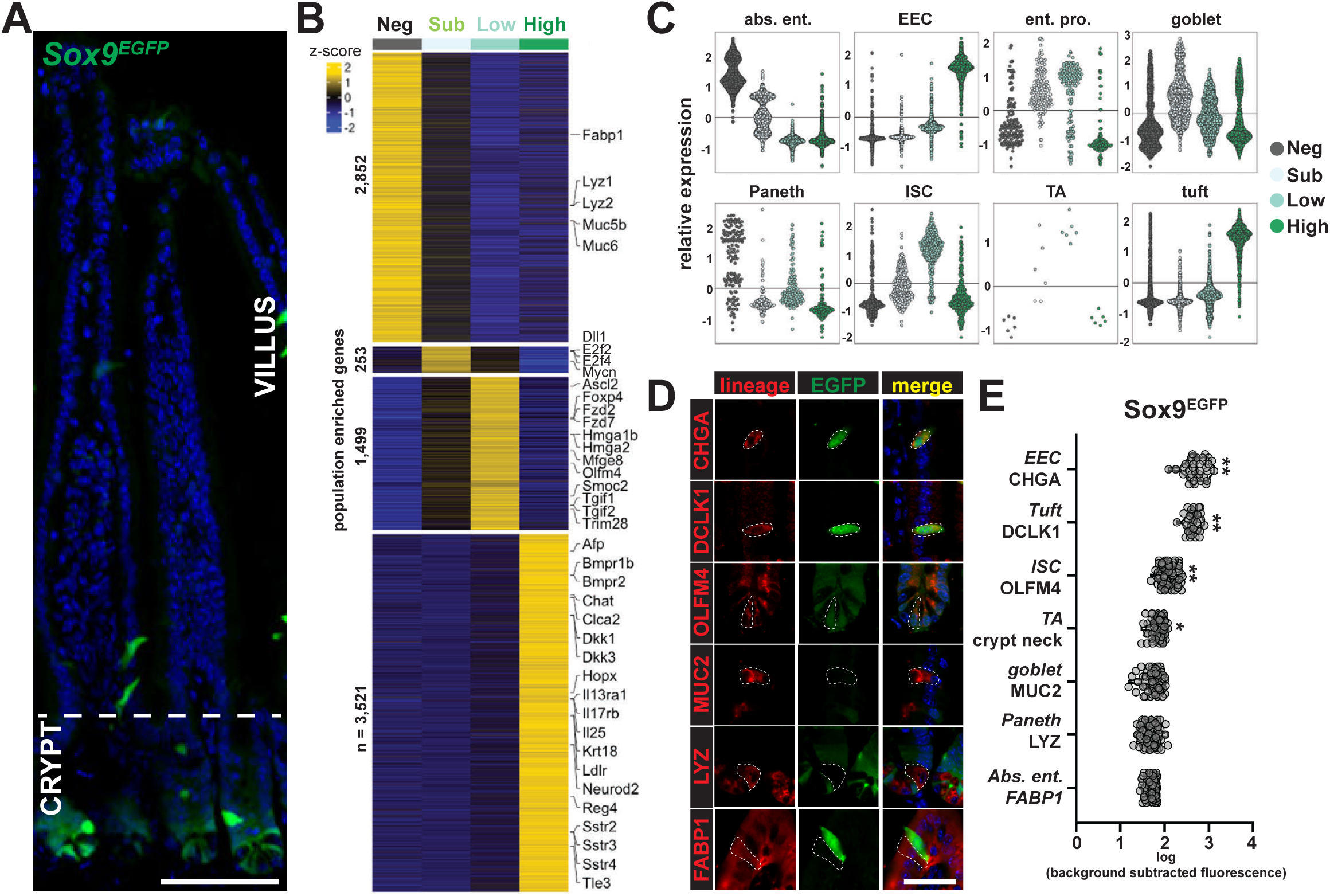
Sox9^EGFP^ populations capture the spectrum of IEC differentiation. (A) Sox9^EGFP^ is expressed at distinct levels in the crypt and as rare “high”-expressing cells in the villi (scale bar represents 100um). (B) Sox9^EGFP^ levels define distinct cell populations bearing unique transcriptomic signatures. (C) Comparison to cell type specific gene expression signatures from scRNA-seq experiments demonstrates relative enrichment of differentiated cell types in *Sox9*^neg^, progenitors in *Sox9*^sub^, ISCs in *Sox9*^low^, and EECs and tuft cells in *Sox9*^high^ (each point represents the expression of one gene). (D) Co-localization between cell type specific protein markers and Sox9^EGFP^ by immunofluorescence and (E) quantification of EGFP signal confirms these relationships (scale bar in D represents 25um; each point in E represents an individual cell; * indicates p < 0.005; ** indicates p < 0.0001).

We then compared gene expression levels from each Sox9^EGFP^ population against gene sets that define distinct IEC subtypes in previously published scRNA-seq data (Fig. 1C) (Haber et al., 2017). Consistent with our prior characterization of Sox9^EGFP^ populations, genes that define absorptive enterocytes and Paneth cells were enriched in *Sox9*^neg^, TA and enterocyte progenitors (EP) in *Sox9*^sublow^, *Lgr5*-positive ISCs in *Sox9*^low^, and EECs and tuft cells in *Sox9*^high^. Similar trends of enrichment between Sox9^EGFP^ populations and IEC-specific gene sets were observed in comparisons to a second, independently-generated scRNA-seq data set (Fig. S1D) (Yan et al., 2017). Interestingly, a portion of goblet cell genes were enriched in *Sox9*^sublow^ TAs, which may reflect lineage “priming” of post-mitotic gene biomarkers in progenitor populations, consistent with previous reports (Kim et al., 2016b).

Finally, we validated correlation between Sox9^EGFP^ levels and protein biomarkers of IEC populations by semi-quantitative confocal microscopy, which demonstrated that cells identified by cell type-specific markers express EGFP levels predicted by RNA-seq (Fig. 1D&E). Importantly, MUC2+ goblet cells were found in the *Sox9*^neg^ population, despite enrichment of some goblet cell genes in *Sox9*^subow^ TAs. Collectively, our RNA-seq data support the distinct identities of Sox9^EGFP^ populations, define the transcriptional output of each population, and establish the Sox9^EGFP^ model as a viable, single-biomarker approach for isolating ISCs, progenitors, and post-mitotic cells for studies of chromatin dynamics.

### Open chromatin regions are dynamic across *Sox9* populations

Next, we sought to determine the contribution of chromatin dynamics to the unique transcriptomic identities of Sox9^EGFP^ populations. We mapped open chromatin regions (OCRs) in *Sox9*^neg^, *Sox9*^sublow^, *Sox9*^low^, and *Sox9*^high^ by Omni-ATAC-seq (Corces et al., 2017). 81,919 high confidence OCRs were identified across all four populations (Fig. S2A and Supplemental Table 3). We observed a strong correlation between biological replicates within each Sox9^EGFP^ population, and distinct clustering of *Sox9*^neg^, *Sox9*^sublow^, *Sox9*^low^, and *Sox9*^high^ by OCRs (Fig 2A). *Sox9*^sublow^ TAs and *Sox9*^low^ ISC populations were more similar by Pearson correlation than *Sox9*^neg^ and *Sox9*^high^ population, consistent with their identity as proliferative, undifferentiated populations (Fig. 2A). We classified peaks found in all four Sox9^EGFP^ populations as “shared” and compared shared peaks to peaks found in single populations relative to genomic features. A majority of peaks present in a single population were found in introns or intergenic regions, while promoters represented a greater relative proportion of shared peaks (Fig 2B). Site-specific analysis of genome browser tracks confirmed dynamic changes in OCRs, with open chromatin found in loci encoding genes associated with function in Sox9^EGFP^ subpopulations. *Lct*, which is expressed in absorptive enterocytes was enriched for OCRs specific in *Sox9*^neg^ (Fig. 2C), with a similar correlation between expression and chromatin status observed for *Neurod1* in *Sox9*^high^ (Fig. 2D) and *Olfm4* in *Sox9*^low^ (Fig. 2E). Some intragenic OCRs were found in Sox9^EGFP^ populations that did not highly express the corresponding gene, as observed for Olfm4-associated OCRs in *Sox9*^sublow^ cells (Fig. 2E). This could either reflect chromatin “priming” to facilitate activation of the corresponding gene under stimulus, or the presence of an enhancer with impacts on a distal gene.

**Figure 2:**
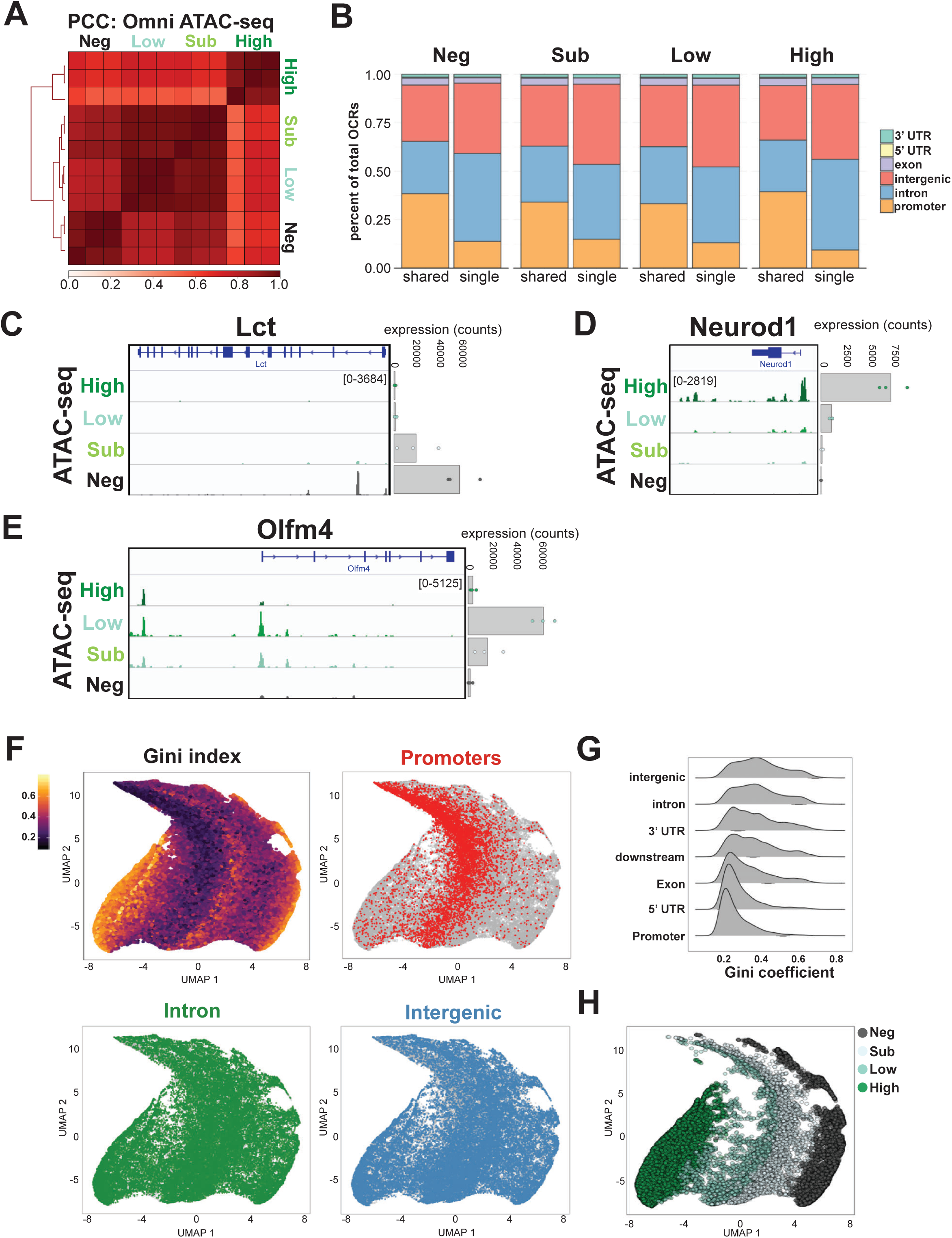
IEC populations have unique and dynamic OCR signatures. (A) Sox9^EGFP^ populations cluster reproducibly by Pearson correlation of OCRs (n = 3 biological replicates per population). (B) Intergenic regions and introns represent the majority of OCRs present in single populations, while promoters represent a larger portion of OCRs shared across all four populations. Representative browser tracks demonstrate relationships between OCR activity and gene expression for (C) *Sox9*^neg^ associated *Lct*, (D) *Sox9*^high^ associated *Neurod1*, and (E) *Sox9*^low^ associated *Olfm4*. (F) UMAP dimensionality reduction and Gini index analysis of OCRs reveals broadly-permissive and dynamic chromatin and predictable clustering by genome feature, including broadly-permissive promoters and dynamic introns and intergenic loci (n = 81,919 OCRs). (G) Quantitative ranking of genome features by Gini index confirms that the highest degree of chromatin dynamics are associated with introns and intergenic regions, but demonstrates dynamic loci in all features, including promoters. (H) UMAP visualization of dynamic OCRs (Gini ≥ 0.4) by highest accessibility demonstrates distinct clustering by Sox9^EGFP^ population.

A limitation of peak overlap based methods for assessing chromatin dynamics in a multi-population experiment is how to classify peaks that are found in 2 or more populations, but not in all populations. In order to more quantitatively assess chromatin dynamics, we analyzed OCR variability across Sox9^EGFP^ populations using Gini index, a statistical measure of inequality that assigns higher values to more variable distributions (Yoshida et al., 2019). We visualized Gini index by Uniform Manifold Approximation and Projection (UMAP), where each point on the plot represents a single OCR and the Gini index indicates the extent of variable accessibility across all four Sox9^EGFP^ populations (Fig. 2F) (McInnes et al., 2018). High Gini index values are associated with dynamic chromatin, while low Gini index values indicate broadly-accessible OCRs (Fig. S2B). As predicted by genome feature analysis of shared and single OCRs, promoters were mostly associated with OCRs bearing low Gini index values, while introns were associated with a greater number of OCRs bearing high Gini index values (Fig. 2F&G). Additionally, promoters were clustered together in the UMAP representation of the data, whereas intronic OCRs were spread throughout the plot, highlighting similarities in accessibility between promoter-associated OCRs. Comparison of Gini values at different genomic features confirmed that the highest degree of dynamic chromatin is found in introns and intergenic regions (Fig. 2G). Despite the overall low diversity of promoters, we identified 738 promoter-associated OCRs with Gini ≥ 0.4 (Fig. 2F&G and Fig. S2; Gini index values for all 81,919 OCRs are listed in Supplemental Table 3). These dynamic promoters were found in all four Sox9^EGFP^ populations (n = 56 *Sox9*^neg^, 97 *Sox9*^sublow^, 216 *Sox9*^low^, 341 *Sox9*^high^) and were associated with genes enriched in their respective population (Fig S2C&D). Applying Gini index values facilitated a more quantitative assessment and visualization of chromatin dynamics, which allowed us to identify OCRs with potential regulatory significance that may have been missed by conventional peak comparisons.

Next, we visualized OCR intensity and distribution by Sox9^EGFP^ population. OCRs for *Sox9*^low^ ISCs and *Sox9*^sublow^ progenitors were distributed across UMAP plots, with overlapping adjacent regions of density (Fig. S2E). *Sox9^neg^*and *Sox9^high^* populations demonstrated highest intensity OCRs at opposite ends of the plot, suggesting that the most significant differences in OCRs are found between populations containing post-mitotic IEC subtypes (Fig. S2E). To focus our analysis, we plotted dynamic OCRs (Gini > 0.4) by Sox9^EGFP^ population in which accessibility was highest. OCRs clustered by Sox9^EGFP^ population, suggesting that many OCRs shared between 2 or more populations are subject to regulatory activity that reaches a maximum in a single cell type (Fig. 2H). Globally, ATAC-seq in Sox9^EGFP^ populations demonstrates that chromatin in IEC differentiation is dynamic, associated with population-defining genes, and exhibits the most significant global changes in introns and intergenic regions in post-mitotic cells.

### Chromatin dynamics are linked to transcription factor networks

We next sought to establish a framework to further examine the relationship between chromatin dynamics and gene expression in Sox9^EGFP^ populations. We reasoned that changes in TF motif usage could be used to identify TFs associated with either broadly-permissive or dynamic IEC chromatin, and further define the *cis*-regulatory landscape relative to potential *trans*-regulatory interactions. chromVAR, which quantifies motif-dependent gain or loss of OCRs, identified altered motif accessibility across Sox9^EGFP^ populations (Schep et al., 2017). Ranking TF motifs by variability in Sox9^EGFP^ populations revealed motifs associated with broadly-permissive chromatin (n = 406 motifs, chromVAR value < 5) as well as dynamic chromatin (n = 195 motifs, chromVAR value ≥ 5) (Fig 3A and Supplemental Table 4).

**Figure 3:**
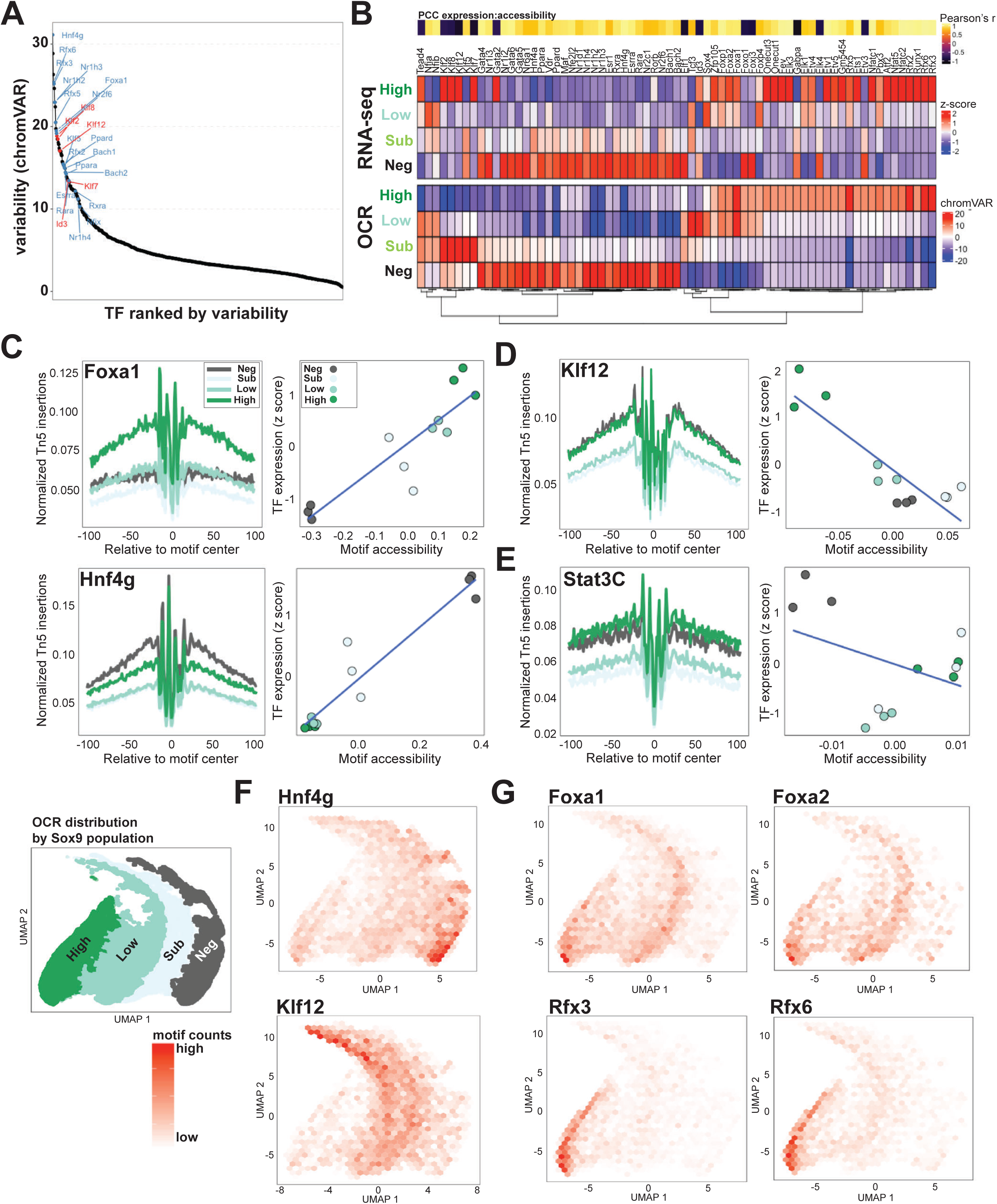
Identification of regulatory networks by OCR dynamics. (A) Analysis of motif usage in Sox9^EGFP^ populations identifies TFs associated with dynamic (n = 195, chromVAR ≥ 5) and broadly-permissive (406 motifs, chromVAR value < 5) OCRs. (B) A subset of 69 TFs demonstrate significant correlation with dynamic motif usage. TFs were selected based on a significant association between accessibility and expression, chromVAR > 5, and TF expression higher than the 25^th^ percentile of all genes. (C-E) Tn5 insertion relative to motif center (left) and relationships between TF expression and motif accessibility (right) (C) A majority of gene expression and motif relationships are positively correlated, including *Foxa1* and *Hnf4g*. (D) Negative correlations are observed for transcriptional repressors including *Klf12*. (E) TFs lacking correlation between motif accessibility and expression, like *Stat3*, are associated with broadly-permissive chromatin. (F) Mapping motif counts with respect to UMAP space plots of Sox9^EGFP^ population specific OCRs reveals distributions predicted by gene expression/motif accessibility relationships (n loci = 12,117 *Hnf4g*; 7,877 *Klf12*). (G) Motifs for *Foxa1/2* and *Rfx3/6* are differentially distributed within *Sox9*^high^ OCRs, suggesting regulatory roles in subpopulations of the same parent Sox9^EGFP^ population (n loci = 7,472 *Foxa1*; 3,302 *Foxa2*; 3,680 *Rfx3*; 4,862 *Rfx6*).

To further explore relationships between TF expression and chromatin dynamics, we analyzed correlation between these two readouts and generated a subset of TFs demonstrating: (1) significant correlation between gene expression and chromatin accessibility over motifs and (2) chromVAR values > 5 (Fig 3B). Increased chromatin accessibility over a given motif was mostly associated with increased expression of the corresponding TF. This category included known regulators of IEC differentiation, which validated the computational approach. *Foxa1*, which drives EEC differentiation, is most highly expressed with the greatest degree of motif accessibility in *Sox9*^high^ cells and *Hnf4g*, which drives absorptive differentiation, is most highly expressed with the greatest degree of motif accessibility in *Sox9*^neg^ cells (Fig. 3C) (Chen et al., 2019; Ye and Kaestner, 2009). We also observed several cases where TF expression and motif accessibility were inversely correlated. *Klf7*, *Klf8*, and *Klf12* were observed to be highly expressed but have the lowest level of motif accessibility in *Sox9*^high^ cells, consistent with known roles for Klf family TFs as repressors (Fig. 3D and Fig. S3A) (McConnell and Yang, 2010). In addition to Klf TFs, we identified other TFs with no established role in ISC differentiation, including nuclear receptor superfamily genes in *Sox9*^neg^, and Ets family genes in *Sox9*^high^ (Fig. 3B). Targeted examination of expression and accessibility relationships for known ISC (Fig. S3B) and early progenitor (Fig. S3C) TFs revealed a mix of expected and unexpected relationships. *Sox9* and *Ascl2* both demonstrated increased expression and motif accessibility in their expected Sox9^EGFP^ populations (Fig. S3B). However, some secretory associated TFs, including *Atoh1* and *Spdef*, demonstrated intermediate accessibility coincident with their highest expression (Fig. S3C). These relationships may suggest competition for or cooperation between multiple TFs at the same motif and require direct detection of TF binding in order to fully understand TF-chromatin relationships.

Finally we visualized TF motif usage in OCRs relative to Sox9^EGFP^ population by UMAP to determine if OCRs enriched for specific motifs formed clusters. Motif accessibility correlated with populations as predicted by accessibility and TF expression relationships and revealed differential distribution of TF motif usage within Sox9^EGFP^ population clusters (Fig. 3F, G). *Foxa1* and *Foxa2* demonstrated similar distribution within *Sox9*^high^ associated OCRs, while *Rfx3* and *Rfx6* motifs were also enriched within the *Sox9*^high^ cluster, but with unique distribution (Fig. 3G). These data suggest the existence of multiple *cis*-regulatory landscapes within each population and are supported by differential regulation of EEC subpopulations by *Foxa1/2* (L- and D-cells) versus *Rfx3/6* (K-cells) (Suzuki et al., 2013; Ye and Kaestner, 2009). By analyzing OCRs in Sox9^EGFP^ populations relative to associated TF motifs, our data infer *trans-*regulatory interactions with dynamic chromatin that are confirmed by identification of TFs with known roles in IEC differentiation. Furthermore, the TF-motif dependent variability in OCR dynamics suggests that different *trans*-regulatory pathways may be associated with either broadly-permissive or dynamic chromatin.

### Chromatin dynamics are predictive of ISC expansion in Id3 null intestines

To test if our computational analyses were robust in identifying novel ISC regulatory factors, we focused on *Id3*, which demonstrates an inverse correlation between TF expression and motif accessibility (Fig. 4A). *Id3* is a known transcriptional repressor with functional significance in colon cancer, but its role in ISC biology remains unknown (Benezra et al., 1990; O’Brien et al., 2012). We acquired Id3 null mice, which carry a red fluorescent protein (RFP) inserted in the first exon of Id3, resulting in loss of expression (McMahon et al., 2008). RFP was observed in the villus of Id3^RFP/RFP^ jejunum, consistent with *Id3* expression in differentiated *Sox9*^neg^ and *Sox9*^high^ populations by RNA-seq (Fig. 4B). Crypts in Id3^RFP/RFP^ intestines were larger compared to wild-type controls (Fig. 4C) and the number of ISCs was significantly increased, based on immunofluorescence for OLFM4, a specific ISC marker (Fig. 4D) (Schuijers et al., 2014). Additionally, we found that OLFM4+ cells extended further up the crypt axis (Fig. 4E). We observed no change in the percent of KI67+ proliferating cells in Id3^RFP/RFP^ crypts, suggesting that increased proliferation does not drive the observed increase in ISCs (Fig. 4F). Collectively, our data demonstrate an expansion in ISC numbers in Id3^RFP/RFP^ knockout mice, as predicted by *Id3* expression and motif dynamics in Sox9^EGFP^ populations. The inverse relationship between *Id3* expression and motif accessibility, combined with the observed phenotype, suggests that *Id3* is involved in the repression of ISC-specific OCRs involved in maintaining the stem cell state.

**Figure 4:**
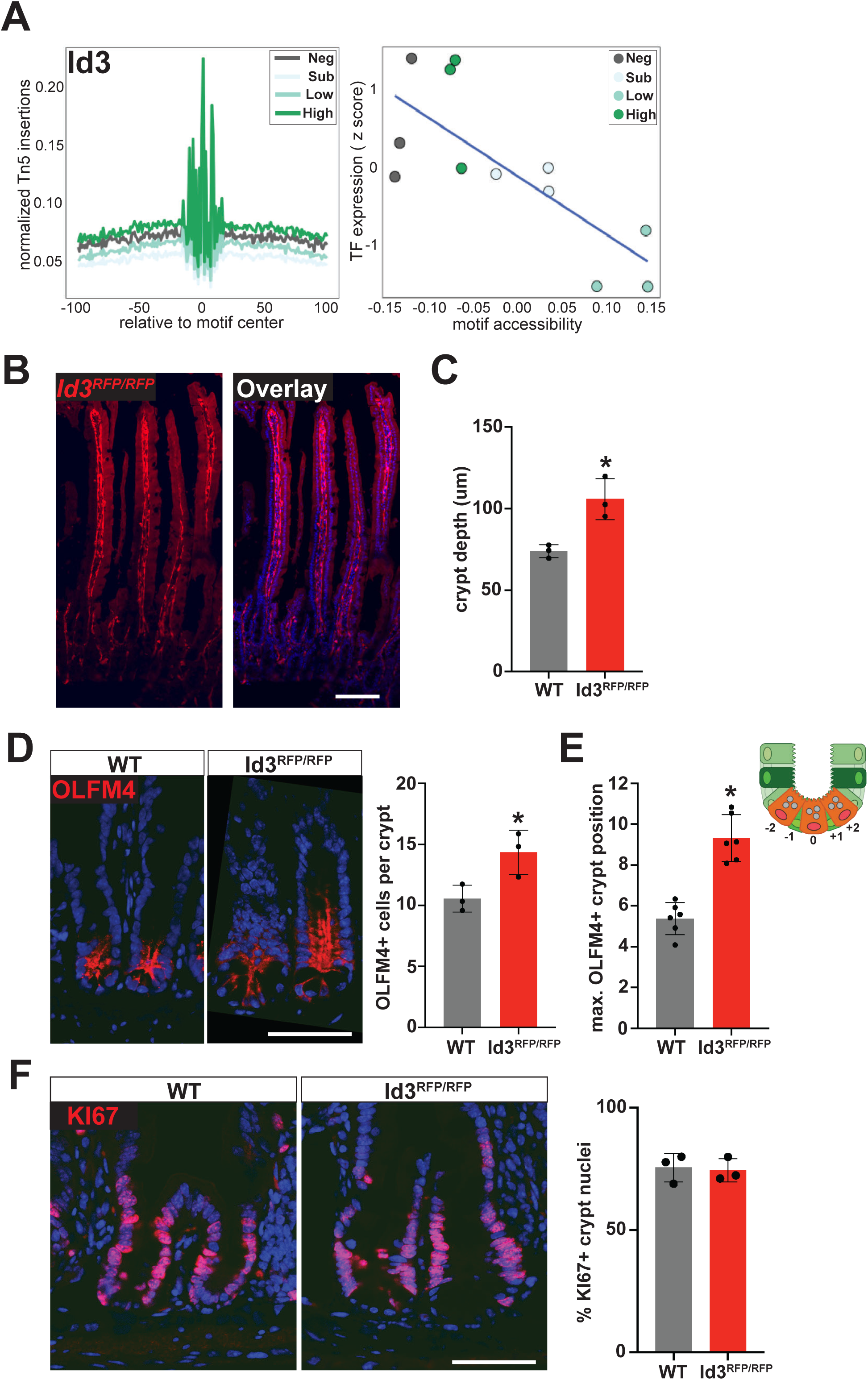
Loss of Id3 results in increased ISC numbers. (A) Tn5 insertion frequency relative to motif center (left) and expression and motif accessibility relationships (right) suggest a repressive role for *Id3* in ISC (*Sox9*^low^) associated OCRs (n = 5,664 loci). (B) Id3^RFP/RFP^ null mice exhibit RFP expression in villus epithelium, consistent with observed expression of *Id3* in *Sox9*^neg^ and *Sox9*^high^ populations (scale bar represents 100um). (C) Id3^RFP/RFP^ crypts are significantly larger compared to wild-type controls and (D) demonstrate increased OLFM4+ ISC numbers (scale bar represents 50um; * indicates p < 0.05; n = 3 biological replicates, 12 crypts per mouse). (E) OLFM4+ cells are observed significantly further up the crypt-villus axis in Id3^RFP/RFP^ intestines, suggesting delayed differentiation of ISCs (* indicates p < 0.05; n = 3 biological replicates, 12 crypts per mouse). (F) Despite increased OLFM4+ cells and crypt depth, percent KI67+ proliferating cells is unchanged between wild-type and Id3^RFP/RFP^ crypts (scale bar represents 50um; n = 3 biological replicates, 10 crypts per mouse).

### Profiling chromatin features shared between active and facultative ISC populations

The precise genetic regulation of IEC progenitor de-differentiation following epithelial injury remains poorly defined, but is associated with rearrangement of chromatin landscapes in facultative ISCs to more closely resemble active ISC chromatin (Jadhav et al., 2017). Previous studies have demonstrated that a subpopulation of *Sox9*^high^ cells is consistent with label-retaining facultative ISCs (Roche et al., 2015). To assay for common chromatin features between active and facultative ISCs in uninjured intestinal epithelium, we identified OCRs found in *Sox9*^low^ (active) and *Sox9*^high^ (facultative) populations, but excluded from *Sox9*^neg^ and *Sox9*^sublow^ (n = 5,092 of 81,919 peaks). *Sox9*^low/high^ shared OCRs were found primarily in introns and intergenic regions, consistent with dynamic OCRs found across all four Sox9^EGFP^ populations (Fig. 5A). Plotting shared OCRs with respect to UMAP space demonstrated overlap between *Sox9*^low^ and *Sox9*^high^ population distributions (Fig. 5B, compare to Fig. 2H), and shared OCRs exhibited generally higher Gini index values relative to all OCRs (Fig. 5C). To determine if *Sox9*^low/high^ shared OCRs were associated with increased transcriptional activity specific to these populations, we identified the nearest gene to each shared OCR and plotted expression across all Sox9^EGFP^ populations. This gene set was significantly upregulated in *Sox9*^low^ and *Sox9*^high^ populations, further validating our analytical approach to identify shared *cis*-regulatory features with regulatory significance (Fig. 5D and S4A).

**Figure 5:**
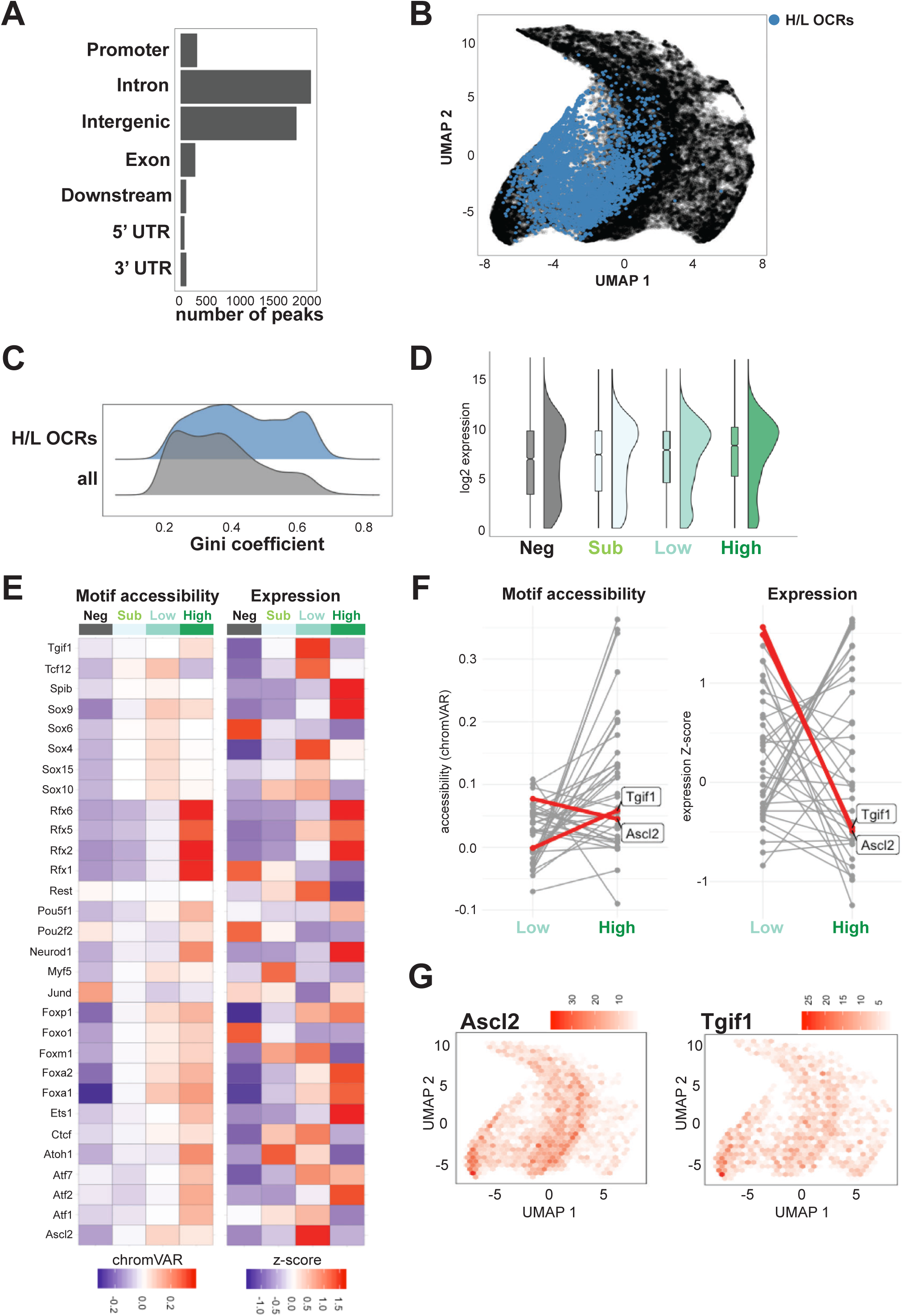
Shared chromatin accessibility in Sox9^low^ and Sox9^high^ populations is associated with enteroendocrine and ISC TFs. (A) *Sox9*^low/high^ shared OCRs are enriched in introns and intergenic regions, consistent with dynamic chromatin characteristics (n = 5,092 OCRs). (B) Shared OCRs localize to overlapping *Sox9*^low^ and *Sox9*^high^ distributions on UMAP and (C) demonstrate increased Gini index values relative to all identified IEC OCRs. (D) Genes nearest to shared OCRs are more highly expressed in *Sox9*^low^ and *Sox9*^high^ populations relative to *Sox9*^neg^ and *Sox9*^sublow^, consistent with functional significance (see Fig. S4A for statistical analysis). (E) 30 TFs demonstrated motif enrichment and expression in *Sox9*^low/high^ shared OCRs, including genes with known roles in ISC self-renewal and differentiation. (F) ISC signature genes *Ascl2* and *Tgif1* exhibit motif accessibility and gene expression patterns consistent with active transcriptional function in *Sox9*^low^ active ISCs and chromatin “priming” in *Sox9*^high^ cells, which contain a known facultative ISC subpopulation. Plots show bias-normalized motif accessibility deviations from chromVAR (left) or z-scored RNA expression (right) in *Sox9*^low^ and *Sox9*^high^ populations for 30 TFs in (E) (G) ISC TF motifs are enriched across OCRs shared or more exclusively associated with *Sox9*^low^ and *Sox9*^high^ cells, suggesting common and distinct regulatory roles in each population (n loci = 5,734 *Ascl2*; 3,281 *Tgif1*).

Next, we sought to leverage the *Sox9*^low/high^ shared OCR data set to identify potential transcriptional regulators of stemness. We identified 30 TF motifs enriched in *Sox9*^low/high^ shared OCRs and compared motif accessibility with gene expression of the corresponding TF (Fig. 5E). This analysis identified two known ISC signature genes, *Ascl2* and *Tgif1* (Munoz et al., 2012). Compellingly, both TFs demonstrated similar motif accessibility between *Sox9*^low^ and *Sox9*^high^ populations, but were expressed significantly higher in *Sox9*^low^ active ISCs (Fig. 5F). *Ascl2* and *Tgif1* motifs were also enriched in shared OCRs as well as OCRs that clustered more strongly with *Sox9*^low^ or *Sox9*^high^ populations (Fig. 5G). These data are consistent with ISC *cis*-regulatory elements being “primed” in both active and facultative ISCs and suggest the presence of shared and unique ASCL2 and TGIF1 binding sites in each population. Conceptually, this would facilitate rapid reactivation of *Ascl2* and *Tgif1* target genes in *Sox9*^high^ facultative ISCs in the setting of damage to the active ISC pool. Similar patterns of accessibility and expression were observed for TFs with no known role in ISC function, including *Atf1* and *Atf7* (Fig. 5E). Future functional assays could be employed to determine if these factors play novel roles in progenitor plasticity and stemness.

We reasoned that, in addition to shared ISC regulatory features, *Sox9*^low^ and *Sox9*^high^ populations would also share OCRs associated with early differentiation from ISCs to EECs. To this end, we identified EEC-associated TFs in our analyses. Unlike ISC TFs, EEC-associated TFs demonstrated substantial increases in both motif accessibility and expression between *Sox9*^low^ and *Sox9*^high^ populations, consistent with the activation of *Sox9*^high^-specific EEC phenotypes (Fig. S5B&C). By examining OCR and gene expression profiles between *Sox9*^low^ and *Sox9*^high^ populations, we characterize chromatin dynamics associated with ISC and EEC TFs and demonstrate proof-of-concept for parsing TF and chromatin networks associated with specific subpopulations from bulk genome/transcriptome analyses.

### 5-hydroxymethylation is dynamic across ISC differentiation

We next sought to examine a specific chromatin modification in Sox9^EGFP^ populations to further assay *cis*-regulatory dynamics. 5hmC is associated with activation of gene expression and can be mapped in small numbers of primary cells using Nano-hmC-Seal (Han et al., 2016). Our transcriptomic data demonstrated that *Tet1*-*3*, which catalyze hydroxymethylation of 5mC, are differentially expressed in Sox9^EGFP^ populations, suggesting the potential for dynamic regulation of 5hmC across ISC differentiation (Fig. S5A). Since 5hmC has been associated with both proximal and distal regulatory elements, we first examined differences in Nano-hmC-Seal signal across Sox9^EGFP^ populations relative to promoters, gene bodies, and distal OCRs (>2kb from TSS) (Fig. 6A). Raw 5hmC signal was enriched over gene bodies (Fig. S5B), but reads were evenly distributed across gene bodies, OCRs, and promoters when normalized for feature length (Fig. S5C). We identified differentially hydroxymethylated regions (DMHRs) in all four Sox9^EGFP^ populations. *Sox9*^neg^ and *Sox9*^high^ populations demonstrated more DMHRs relative to *Sox9*^sublow^ and *Sox9*^low^, consistent with increased 5hmC in differentiated IECs (Fig. 6B) (Kim et al., 2016a). Further, a majority of DHMRs were found over gene bodies in each Sox9^EGFP^ population (Fig. 6B). 5hmC varied across populations with particularly distinct hydroxymethylated promoters and gene bodies in *Sox9*^neg^ and *Sox9*^high^ populations, while OCRs demonstrated more shared or potentially progressive hydroxymethylation across all Sox9^EGFP^ populations (Fig. 6C).

**Figure 6:**
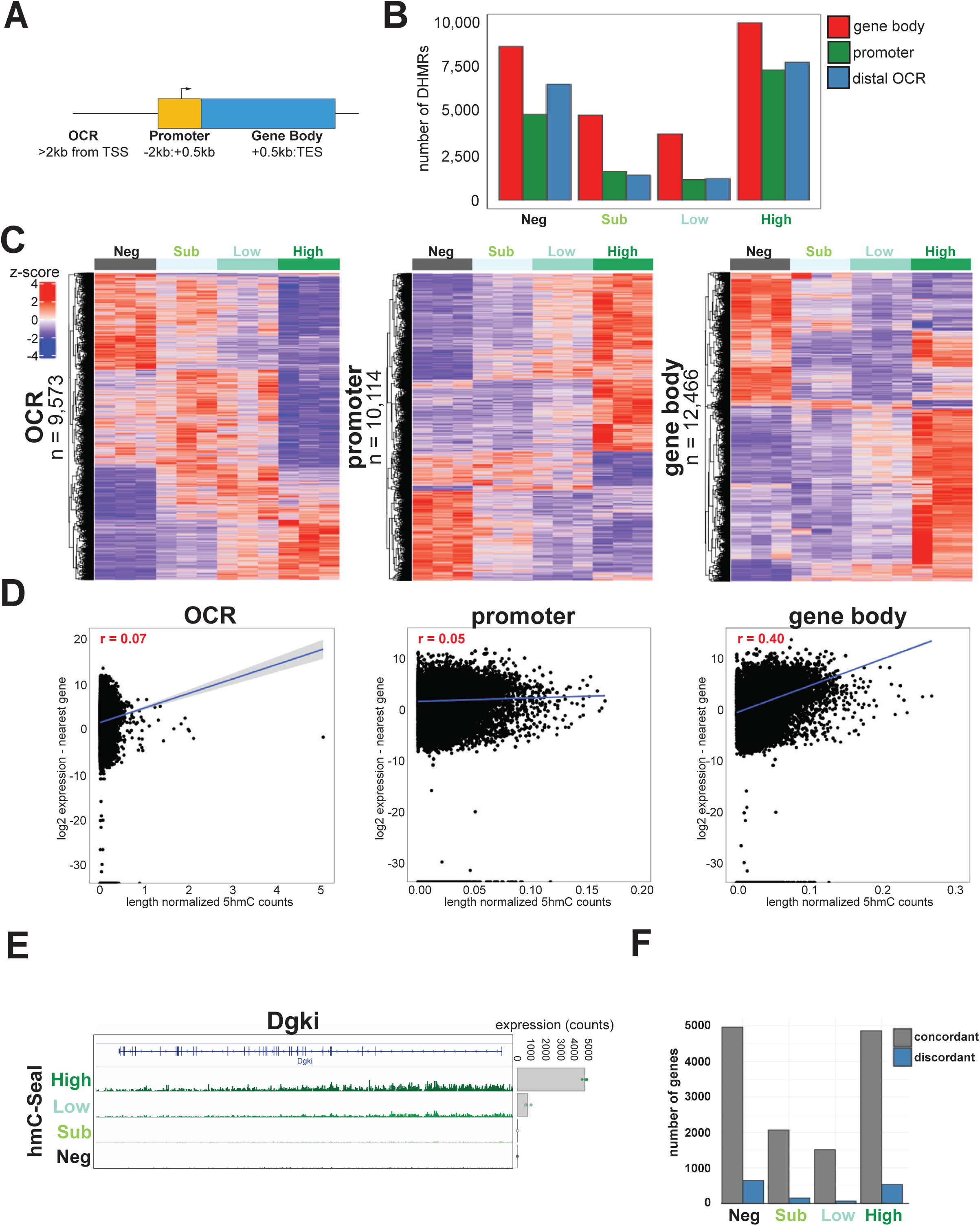
5hmC is associated with OCR dynamics in putative distal regulatory regions. (A) Selection criteria for the three classes of genomic features examined. Read counts from each region were used as input for differential hydroxymethylation analysis. (B) Number of statistically significant DHMRs found in each Sox9^EGFP^ population (adjusted p-value < 0.05). (C) Heatmap of z-scored 5hmC signal at DHMRs from each genomic class. (D) Relationships between 5hmC and gene expression at different genomic classes demonstrate correlation between gene body hydroxymethylation and gene expression (Pearson’s r is reported for each association). (E) Browser tracks of 5hmC signal at *Dgki* show that intragenic 5hmC is dynamic across Sox9^EGFP^ populations and associated with changes in gene expression. (F) A majority of genes demonstrate concordant relationships between 5hmC and expression.

To infer the regulatory significance of 5hmC distribution by genomic element, we compared 5hmC levels to expression levels of genes located nearest to 5hmC-enriched loci. While OCRs and promoters demonstrated little correlation with gene expression, gene body 5hmC correlated positively with gene expression (Fig. 6D). Interestingly, we also noted a positive correlation between promoter and gene body 5hmC levels, suggesting that despite relationships between promoter and genic 5hmC, the latter is more predictive of gene expression levels (Fig. S5D). We noted the same moderate positive correlation between expression and genic 5hmC in all Sox9^EGFP^ populations, with *Sox9*^high^ and *Sox9*^neg^ cells displaying the strongest relationship, consistent with increased DHMRs in these populations and previous reports of 5hmC enrichment in villus epithelium (Fig. S5E) (Kim et al., 2016a). Accordingly, genome browser tracks revealed broad, dynamic domains of 5hmC across gene bodies that correlate with gene expression between Sox9^EGFP^ populations (Fig. 6E). Since genic 5hmC has been linked to increased transcription in previous studies, we compared fold change in gene expression to fold change of gene body 5hmC in each Sox9^EGFP^ population (Han et al., 2016; Tsagaratou et al., 2014; Wu et al., 2011). We found that the vast majority of genes in each Sox9^EGFP^ population were concordantly regulated relative to changes in genic 5hmC (gene expression increasing with genic 5hmC) (Fig. 6F). Together, our Nano-hmC-seal data demonstrate that 5hmC is differentially enriched between Sox9^EGFP^ populations across the genome, but that gene body 5hmC is most predictive of gene expression levels.

### Genic 5-hydroxymethylation correlates with absorptive gene expression

To determine if 5hmC dynamics correlate with predictable patterns in ISC differentiation, we assessed the enrichment of 5hmC signal over defined marker genes for cell types identified by scRNA-seq (Haber et al., 2017). Consistent with previous reports, this analysis showed that 5hmC is depleted near the TSS, with high levels of the mark found throughout the gene body (Fig. 7A and S6A) (Han et al., 2016). We observed predictable relationships between enrichment of 5hmC in gene bodies associated with specific IEC subpopulations and their corresponding Sox9^EGFP^ population. For example, *Sox9*^neg^ cells were enriched for high 5hmC signal over genes associated with absorptive enterocytes, while *Sox9*^high^ cells demonstrated modest enrichment of 5hmC over genes associated with tuft cells (Fig. 7A and S6A). Most cell type specific marker gene sets were devoid of 5hmC enrichment associated with any one Sox9^EGFP^ population. This suggests that 5hmC may have regulatory significance in some IEC populations, while being less important for cell type specific gene expression in others.

**Figure 7:**
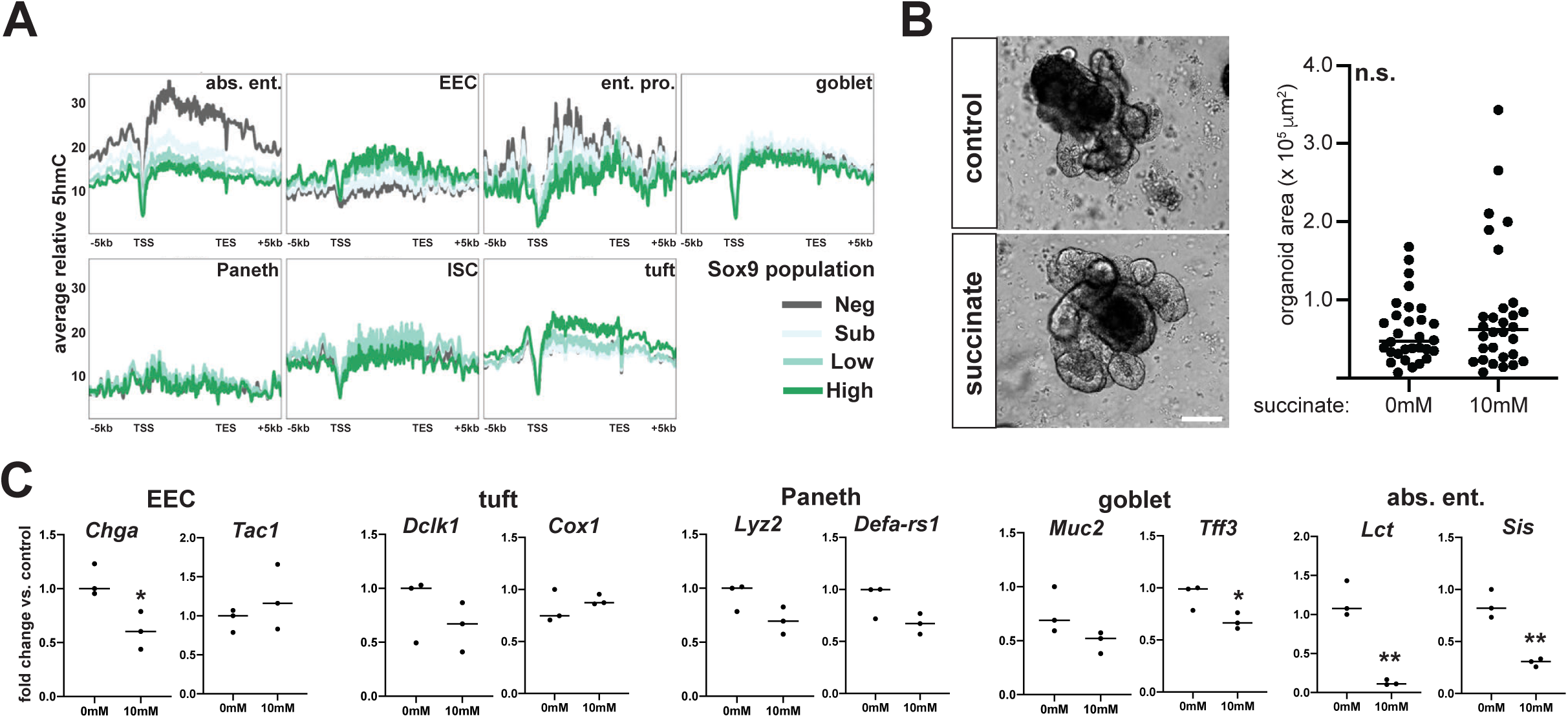
5hmC dynamics over gene bodies are predictive of a role in absorptive gene expression. (A) Enrichment of 5hmC over scRNA-seq defined IEC gene sets reveals increased signal in absorptive enterocyte genes in the *Sox9*^neg^ population and tuft cell genes in the *Sox9*^high^ population. (B) Organoids treated with dimethyl succinate to suppress global 5hmC levels do not demonstrate changes in organoid size (n = 30 organoids per 0mM and 10mM dimethyl succinate groups, from 3 biological replicates; scale bar represents 100um). (C) Three days of succinate treatment results in a pronounced downregulation of absorptive enterocyte markers *Sis* and *Lct*, as well as decreased expression of *Chga* and *Tff3* (n = 3 0mM and 10mM dimethyl succinate; * indicates p < 0.05, ** indicates p < 0.005).

To functionally test the role of 5hmC, we sought to modulate enzymatic activity of TETs in intestinal organoid cultures. TET1-3 are expressed in an overlapping manner in IECs, and site-specific roles, potential redundancy, and non-enzymatic function complicate 5hmC functional analyses relying on TET knockdown. Since all TET enzymatic activity is sensitive to intracellular succinate/alpha-ketoglutarate ratios, we treated organoids with cell-permeable succinate to inhibit oxidation of 5mC to 5hmC, as previously applied in ES cells (Carey et al., 2015). Dot blot analysis revealed a significant reduction in global 5hmC levels in organoids treated with 10mM succinate for 5 days *in vitro* (Fig. S6B&C). Succinate-treated organoids appeared normal following treatment, with no obvious morphological defects or changes in size (Fig. 7B). To assess lineage allocation, we conducted RT-qPCR for IEC subtype-specific markers in control and succinate-treated organoids. We observed a significant and reproducible downregulation of absorptive enterocyte specific genes, *Lct* and *Sis*, at both 3 and 5 days of succinate treatment (Fig. 7C and Fig. S6D). While *Chga* and *Tff3* demonstrated slight downregulation with 3 days of treatment, this effect was not observed at 5 days, owing perhaps to increasing organoid heterogeneity and variability in gene expression between biological replicates over time. Succinate treatment had no impact on ISC gene expression (Fig. S6E). Our organoid experiments suggest that Nano-hmC-Seal assays demonstrating the greatest degree of 5hmC enrichment over absorptive enterocyte genes are predictive of the sensitivity of these genes to decreased global 5hmC.

## DISCUSSION

Prior studies have arrived at conflicting conclusions regarding the extent to which chromatin in IECs is broadly-permissive (Kaaij et al., 2013; Kim et al., 2014) or dynamic, with cell-type specific patterns in accessibility and chromatin modifications (Jadhav et al., 2017; Kazakevych et al., 2017; Lindeboom et al., 2018). Broadly-permissive chromatin has been proposed as the genomic mechanism that facilitates progenitor de-differentiation during intestinal regeneration (Kim et al., 2014). However, focused studies of a secretory progenitor subpopulation have demonstrated significant chromatin reorganization during this same process, suggesting that chromatin must also undergo a significant degree of reorganization during normal differentiation (Jadhav et al., 2017). In an attempt to resolve these apparently conflicting results, we “mapped” OCRs and 5hmC in IEC populations identified by a Sox9^EGFP^ transgene. While the Sox9^EGFP^ model does not isolate “pure” IEC subpopulations, our transcriptomic analyses demonstrate that distinct EGFP expression levels are capable of reproducibly defining ISCs, TAs, and differentiated populations, capturing the longitudinal spectrum of lineage specification and facilitating “tissue wide” chromatin profiling in an unperturbed system. This study establishes the Sox9^EGFP^ transgene as tool for studying chromatin regulation in multiple IEC subpopulations, including intermediate progenitors, without requiring genetic modulation to enrich for different lineages.

Defining the relative extent of inter-population chromatin differences as “broadly-permissive” or “dynamic” remains a significant conceptual challenge. Logically, all cell types within a single tissue can be expected to share a large number of broadly-permissive *cis*-regulatory elements foundational to tissue identity. It is also expected that some degree of *cis*-regulatory elements define requirements for cellular subpopulation identity, and that these would appear more dynamic across populations. To dissect broadly-permissive and dynamic elements, we applied Gini index as a quantitative approach for objectively defining changes in OCRs. The Gini index was developed for economic dispersion analysis, but was recently applied to quantify chromatin dynamics in the hematopoietic system (Yoshida et al., 2019). Our data affirm Gini index as a valuable approach for quantifying chromatin dynamics in studies involving 3 or more groups, where pairwise analyses may be insufficient. We found that a majority of OCRs with relatively high Gini index values were present in intronic or intergenic regions, consistent with the positioning of enhancer elements and genome features associated with highly dynamic chromatin in the hematopoietic system (Yoshida et al., 2019). Functional association between specific enhancers and their associated genes would help further define the significance of these *cis-*regulatory elements in establishing and maintaining IEC cell fates.

To further characterize chromatin in Sox9^EGFP^ populations, we combined analysis of (1) OCR activity, (2) TF motifs found within OCRs, and (3) corresponding TF expression to identify transcriptional networks and their associated *cis*-regulatory landscape. Most identified TF motifs (n = 406 motifs) were associated with OCRs that exhibited little variation in accessibility across ISC differentiation, consistent with a model of broadly-permissive chromatin, where IEC chromatin is “primed” to respond to diverse transcriptional regulatory signals (Kim et al., 2014). However, we identified an additional 195 motifs that were associated with dynamic chromatin accessibility, including known and potentially novel regulators of specific IEC lineage differentiation. We validated our computational approach by demonstrating a previously uncharacterized expansion of ISCs in Id3 knockout mice, which was predicted by loss of ID3 motif accessibility found in *Sox9*^low^ ISCs that correlated with increased *Id3* expression in differentiated IECs. A notable limitation of this approach is the reliance on predicted TF motifs, which may overlap between multiple TFs. In the case of ID3, which does not directly bind DNA, the repressive effect on chromatin accessibility may be due to interaction with multiple distinct TFs (Benezra et al., 1990). Direct mapping of TF-DNA interactions in Sox9^EGFP^ populations will be required to further understand the interaction between TF networks and broadly-permissive or dynamic IEC chromatin. Our data provide a comprehensive resource of expression and chromatin accessibility that can be used as a foundation for functional and mechanistic studies that uncover new regulators of IEC function.

To address the limitation of Sox9^EGFP^ populations containing multiple subpopulations of different IECs, we demonstrated that “multi-omic” analyses can be used to parse useful regulatory information from bulk data sets. Specifically, we applied quantitative analyses to identify shared OCRs, TF motifs, and corresponding TF expression between *Sox9*^low^ and *Sox9*^high^ populations. This approach allowed us to characterize expression and motif accessibility dynamics for ISC-associated and EEC-associated TFs. In line with proposed mechanisms for progenitor plasticity, we found that ISC-associated TFs *Ascl2* and *Tgif1* exhibit characteristics consistent with broadly-permissive chromatin, with variable expression but relatively invariable accessibility between *Sox9*^low^ and *Sox9*^high^ populations. In contrast, EEC-associated TFs are associated with both highly variable expression and motif accessibility, suggesting different *cis*- and *trans*-regulatory mechanisms underlying cellular processes of plasticity and lineage specification.

To understand a specific chromatin modification in ISC differentiation, we examined 5hmC dynamics. In agreement with previous studies in embryonic stem cells and the hematopoietic system, we observed enrichment of 5hmC in putative distal enhancers as well as promoters and gene bodies, but found that only genic 5hmC correlates with gene expression level (Han et al., 2016; Tsagaratou et al., 2014; Wu et al., 2011). To infer the functional significance of this correlation, we combined Sox9^EGFP^ population identities with gene sets identified in published scRNA-seq to plot enrichment of 5hmC over cell-type specific gene sets. This revealed increased hydroxymethylation over absorptive enterocyte genes, which we functionally confirmed by observed downregulation of *Lct* and *Sis* following succinate treatment of intestinal organoids. While succinate treatment results in global reduction of 5hmC, it has many other effects, including inhibition of other alpha-ketoglutarate-dependent chromatin modifying enzymes (Carey et al., 2015). Though outside of the scope of the present study, our organoid experiments raise questions about the individual and collective role of TET enzymes and 5hmC in ISC differentiation. While no current mechanism explains the link between genic 5hmC and increased gene expression, it has been speculated that this is secondary to TET2/3 association with RNA Pol II via SET/COMPASS (Deplus et al., 2013). Therefore, it remains unknown whether the observed downregulation of 5hmC-enriched absorptive genes in our experiments is a direct consequence of, or a correlation with, globally reduced 5hmC. Previous studies have shown that *Tet1* is required for proper ISC numbers (Kim et al., 2016a), and further work dissecting the 5hmC-dependent and –independent functions of TET enzymes is needed to fully understand the role of 5hmC in ISC biology.

Large-scale chromatin mapping projects, including the Epigenome Roadmap, are critical for expanding our understanding of the functional genome, but often focus on whole tissues at the expense of cell-level resolution (Roadmap Epigenomics et al., 2015). Next generation studies require higher resolution approaches to avoid population averaging effects and fully understand the regulatory significance of chromatin landscapes. Our study provides a chromatin “roadmap” of a continuously renewing adult tissue that will facilitate further mechanistic dissection of the chromatin modifying enzymes and TF networks involved in establishing functional genomes in IECs. Our quantitative analyses demonstrate that the IEC chromatin landscape can be both broadly-permissive and highly dynamic, and that the extent of chromatin changes in ISC differentiation can be classified relative to predicted TF networks.

## METHODS

### Mice

*Sox9^EGFP^* mice were maintained as heterozygotes on the C57Bl/6 background. Id3^RFP/RFP^ mice were obtained from the Zhuang lab at Duke University and were also maintained on the C57Bl/6 background. Wild-type C57Bl/6 mice were used as controls in Id3 knockout experiments All genomics experiments were carried out on adult males between 8-16 weeks of age. Id3 knockout experiments were carried out on adult male and female mice between 8-20 weeks of age, and control and Id3^RFP/RFP^ mice were age and sex-matched. Mice were co-housed, littermates were used when possible, and age-matched non-littermates when not. All mice received Tekland Global Soy Protein-Free Extruded Rodent Diet (Envigo, 2020SX) and water *ad libitum*. Phenotyping for *Sox9^EGFP^* was carried out by observation of EGFP expression in hair follicles of tail clippings, as previously described (Formeister et al., 2009). The Institutional Animal Care and Use Committee (IACUC) of the University of North Carolina at Chapel Hill reviewed and approved all protocols.

### Intestinal epithelial dissociation and FACS

Full length intestine (duodenum, jejunum, and ileum) was used for all sequencing-based experiments, and jejunum was used for organoid experiments. Intestinal epithelial dissociations were carried out as previously described (Zwarycz et al., 2018). For full length preps, the entire small intestine was dissected and opened longitudinally before being rinsed in 1X DPBS (Gibco) to remove fecal matter. For jejunal preps, the proximal 5cm of intestine was removed and discarded, and the proximal half of the remaining intestine was opened longitudinally and rinsed in 1X DPBS to remove fecal matter. Rinsed intestine was transferred to a 50mL conical containing 10mL of 3mM EDTA (Corning) and incubated for 15min at 4C with gentle nutation (RPM = 40). Intestinal tissue was retrieved and transferred to a glass plate prior to being “brushed” with a pipette tip to remove ∼50% of villi. The tissue was then minced to ∼3-5mM pieces and transferred to a new 50mL conical containing 10mL of 3mM EDTA and incubated for and additional 35min at 4C with gentle nutation (RPM = 40). Tissue fragments were transferred to 10mL DPBS in a third 50mL conical and shaken by hand for 4min to remove crypt-enriched epithelium. To obtain single cells, crypt epithelium was rinsed once with 25mL DPBS then resuspended in 10mL HBSS with 0.3U/mL dispase (Corning) and 10uM Y27632 (Selleck Chem), and incubated at 37C for 12-16min with gentle shaking and observation by light microscopy every 2min until ∼80% of prep consisted of single cells. Cells were rinsed twice with DPBS containing 10% FetalPlex (Gemini Biosciences) to quench dispase activity.

Dissociated single cells were resuspended in 1X Sort Media [Advanced DMEM/F12 (Gibco), N2 (Gibco), B27 (Gibco), 10mM HEPES (Corning), Glutamax (Gibco), Pen/Strep (Gibco), and 10uM Y27632 (Selleck Chem)] and stained with CD31-APC (Biolegend) and CD45-APC (Biolegend) for 45min on ice. Stained cells were rinsed once with Advanced DMEM/F12 and resuspended in 1X Sort Media. 7-AAD (Biolegend) and Annexin V-APC (Biolegend) were added immediately prior to sorting. Cells were sorted on a Sony SH800 FACS instrument with a 100um disposable nozzle “chip”. Gating was carried out as previously described in (Gracz et al., 2018) and shown in Supplemental Figure 1A. Doublet discrimination was visually validated by sorting ∼500 cells directly on to a glass slide and observing by light microscopy.

### Organoid culture and dimethyl succinate treatment

Organoids were cultured from jejunal crypts in Cultrex Type II Growth Factor-Reduced matrix (R&D Biosystems). For 5hmC dot blot experiments, ∼300 crypts were seeded in 30uL Cultrex droplets in 48 well plates and overlaid with 200uL organoid media. For gene expression experiments, ∼100 crypts were seeded in 10uL Cultrex droplets in 96 well plates and overlaid with 100uL organoid media. Media consisted of: Advanced DMEM/F12 (Gibco), N2 (Gibco), B27 (Gibco), 10mM HEPES (Corning), Glutamax (Gibco), Pen/Strep (Gibco), 10% RSPO1-conditioned media (RSPO1 HEK293 cells were kindly provided by Calvin Kuo, Stanford University), 50ng/mL recombinant murine EGF (Gibco), and 100ug/mL recombinant human Noggin (Peprotech). 10uM Y27632 (Selleck Chem) was added at initial plating. Organoids were allowed to establish for 48hr prior to treatment with dimethyl succinate (Sigma). Media and succinate were replaced every 48hr.

### RNA isolation

12,000 cells from each Sox9^EGFP^ population were sorted directly into 500uL of Lysis Buffer (Ambion) and stored at -80C until RNA isolation. Total RNA was prepared using the RNAqueous Micro Kit (Ambion) according to manufacturer instructions and eluted in 15uL molecular-grade H2O (Corning). RNA was treated with Ambion RNAqueous Micro Kit DNase I for 30min at 37C to eliminate gDNA and DNase was inactivated using DNase Inactivation Beads. gDNA-free RNA was transferred to low-binding tubes and stored at -80C until QC and library preparation.

### gDNA isolation for Nano-hmC-Seal-seq and dot blot

For FACS isolated Sox9^EGFP^ populations, cells were collected directly into 250uL 1X sort media, pelleted by centrifugation at 6,000g for 5min, then lysed in 200uL gDNA Lysis Buffer (10mM Tris HCl pH 8.0, 100mM EDTA pH 8.0, 0.5% SDS) with 1mg/mL Proteinase K (Ambion) and incubated at 55C in a water bath overnight. For organoids, growth media was isolated from wells and Cultrex and organoids were lysed in 500uL gDNA lysis buffer with 1mg/mL Proteinase K (Ambion) and incubated at 55C in a water bath overnight. gDNA from both FACS isolated cells and organoids was purified using a Quick DNA MicroPrep kit (Zymo) and assayed for concentration using the Qubit dsDNA High Sensitivity kit (Thermo Fisher) and a Qubit Fluorometer (Thermo Fisher).

### RNA-seq: QC, library prep, and sequencing

RNA from 12,000 cells per Sox9^EGFP^ population was collected for each RNA-seq sample. 5uL of 15uL total from each sample was allocated for RT-qPCR validation. cDNA was prepared as described in “RT-qPCR” and sequential enrichment of *Sox9* mRNA across EGFP-sorted populations was validated by RT-qPCR with Taqman probes against *Sox9* (Life Technologies) using *18S* as an internal control. To validate RNA quality, 1uL of 15uL total from each sample was subjected to Agilent Bioanalyzer analysis using the Total Eukaryotic RNA Pico kit (Agilent). All samples demonstrated significant enrichment of *Sox9* across EGFP population and had RIN values ≥ 8.0. Libraries were prepared using the SMARTer Stranded Total RNA-seq Kit v2 (Clontech) and 8uL of 15uL total RNA per sample, as per manufacturer instructions. Libraries were sequenced on a NextSeq500 (Illumina) with 75bp single-end reads, v2 chemistry. Each library was sequenced to ≥ 2.9 x 10^7^ reads.

### Omni-ATAC-seq: library prep and sequencing

12,000 cells per Sox9^EGFP^ population were collected directly into 250uL 1X sort media, pelleted by centrifugation at 2,000g for 5min, and subjected to Omni-ATAC as previously described in (Corces et al., 2017). Libraries were sequenced on a NextSeq500 (Illumina) with 75bp paired-end reads, v2 chemistry. Each library was sequenced to ≥ 5.0 x 10^7^ reads.

### Nano-hmC-Seal-seq: library prep and sequencing

12,000 cells per Sox9^EGFP^ population were collected via FACS (above). gDNA was isolated as described above (see: *gDNA isolation for Nano-hmC-Seal-seq and dot blot*) and subjected to Nano-hmC-Seal as previously described (Han et al., 2016). Libraries were sequenced on a NextSeq500 (Illumina) with 75bp single-end reads, v2 chemistry. Each library was sequenced to ≥ 2.2 x 10^7^ reads.

### RNA-seq: analysis

Quantification was performed using salmon (version 0.8.2) (Patro et al., 2017) and differential expression testing was performed using DESeq2 (version 1.2.22) with tximport (1.10.1) to summarize transcript data at the gene level in R. Expression values in the Sox9-defined populations were then compared to single cell data from two independent sources (Haber et al., 2017; Yan et al., 2017). Genes for Figure 1C were selected by filtering for genes significantly upregulated in a single population only. Ontology enrichment analysis was performed using clusterProfiler R Package (3.12.0)

### Omni-ATAC-seq: analysis

Paired-end 76 bp reads were trimmed using trim_galore (version 0.4.3), aligned using bowtie2 (version 2.3.4) with -X 2000. Then duplicates were removed using picard (version 2.10.3) and peaks were called with Macs2 (version 2.1.2) –q 0.001 –f BAMPE –keep-dup-all. For visualization, signal tracks were computed using Deeptools (version 2.5.2) bamCoverage – binSize 1 –extendReads –normalizeUsingRPKM –ignoreForNormalization chrM chrX chrY – minFragmentLength 40 –maxFragmentLength 130. Comparison of replicates was performed using bamCompare in the deeptools package. Normalized Tn5 insertions were calculated using pyatac (Schep et al., 2015).

To identify differences in motif usage the R package chromVAR was used (Schep et al., 2017). To compare differences in accessibility at individual sites, UMAP dimensionality reduction was used (McInnes et al., 2018) (R package version 0.2.1.0. https://CRAN.R-project.org/package=umap) and the gini index for each site across the 4 population (3 replicates each) was calculated using the gini function in the edgeR package. Motif counts were identified using the motifmatcher R package, using the default p-value cutoff of 5e-5 and projected into UMAP space. For TFs selected in Fig. 3B, we: (1) applied a filter for overall expression (>25^th^ percentile of all gene expression), (2) required the TF be significantly differentially expressed in at least one Sox9^EGFP^ population, and (3) required that the correlation between motif accessibility and TF expression be significant (adjusted p-value < 0.05). The TFs identified in Fig. 5D were selected based on their enrichment using Homer (findMotifsGenome.pl version 4.8.3) comparing peaks shared between Sox9^low^ and Sox9^high^ with those peaks that were found in Sox9^neg^ and Sox9^sublow^. They were further filtered by the overall expression amount (> 25^th^ percentile of all gene expression).

### Nano-hmC-Seal-seq: analysis

5hmc data were trimmed using trim_galore and aligned to mm9 using bowtie2 (version 2.3.4) – sensitive. Follwing duplicate removal using picard (version 2.10.3), signal tracks were generated using deeptools (version 2.5.2) bamCoverage –binSize 10 –extendReads 150 – normalizeUsingRPKM. Signal over genic regions was calculated using deeptools computeMatrix function and gencode annotations (vM1). Pearson correlation between expression and 5hmc levels was calculated in R using the cor.test function and statistical inferences were adjusted for multiple testing using the p.adjust function. For DHMR analysis, reads in the different genomic bins (Fig. 6A) were counted using the R package featureCounts and then used as input to a DESeq2 analysis as described in the RNA-seq analysis section.

### 5hmC dot blot

gDNA prepared from control and dimethyl succinate-treated organoids was diluted to 400ng per sample in 20uL TE. 40uL 0.1M sodium hydroxide was added and samples were incubated at 95C for 10min in thermal cycler to denature DNA. 60uL ice cold 1M ammonium acetate was added and serial dilutions were made with 60uL sample and 60uL TE. Denatured gDNA was spotted onto Hybond N+ membranes (GE Amersham) using a BioDot SF Microfiltration apparatus (BioRad). Membranes were dried for 5min at 80C, UV crosslinked, and blocked overnight in 5% non-fat dry milk in PBS. Membranes were incubated in anti-5-hydroxymethylcytidine (Active Motif, mouse monoclonal, 59.1, 0.4ug/mL) for 3hr at RT in 5% non-fat dry milk in PBS, then rinsed three times in PBS. Secondary detection was carried out by incubating membranes in Gt-anti-Ms HRP (Jackson Immunoresearch, 1:5000) for 3hr at RT in 5% non-fat dry milk in PBS. Membranes were rinsed three times in PBS, HRP was detected using Clarity ECL (BioRad), and blots were imaged using a Protein Simple gel box/imaging station. Following 5hmC detection, blots were rinsed three times in PBS and incubated for 10min at RT in 0.02% methylene blue (Sigma) in 0.3M sodium acetate, pH 5.2 to detect total gDNA. Methylene blue-stained blots were rinsed 5 times in molecular grade H2O and imaged using a Protein Simple gel box/imaging station. Signal intensity for 5hmC was measured for 100ng dots in ImageJ and changes in signal for succinate-treated organoids was calculated as percent change relative to the untreated control for each biological replicate (n = 4 biological replicates).

### RT-qPCR

cDNA was prepared using iScript cDNA Synthesis Kit (BioRad) and diluted 1:10 with molecular grade H2O (Corning) prior to use in RT-qPCR assays with SsoAdvanced Universal Master Mix (BioRad). Taqman primers (Life Tech) were used to detect: *Ascl2* (Mm01268891_g1), *Chga* (Mm00514341_m1), *Cox1* (Mm04225243_g1), *Defa-rs1* (Mm00655850_m1), *Dclk1* (Mm00444950_m1), *Lct* (Mm01285112_m1), *Lgr5* (Mm00438890_m1) *Lyz2* (Mm00727183_s1), *Muc2* (Mm00545872_m1), *Sis* (Mm01210305_m1), *Sox9* (Mm00448840_m1), *Tac1* (Mm00436880_m1), *Tff3* (Mm00495590_m1). Fold change was calculated using ΔΔC_T_ with *18S* (Hs99999901_s1) as the internal reference (Pfaffl, 2001).

### Immunofluorescence and image quantification

*Sox9^EGFP^*, Id3^RFP/RFP^, or wild-type C57Bl/6 control jejunum was dissected, fixed overnight in 4% PFA, then transferred to 30% sucrose for an additional 16-18 hours prior to embedding in Tissue-Tek OCT (Sakura) and frozen at -80C. Immunofluorescence for IEC lineage markers was carried out on 10um sections. Tissue was permeabilized with 0.3% Triton-X 100 (Sigma) in PBS for 20 minutes at room temperature, followed by blocking in 1x Animal-Free Block (Cell Signaling Technologies) for 40 minutes. Primary antibodies (Supplemental Table 6) were diluted at specified concentrations in 1x Animal-Free Block and incubated for 2hr at RT. Secondary antibodies were diluted 1:500 in 1x Animal-Free Block and incubated for 45min at RT. Nuclear counterstain was carried out by bisbenzimide (Sigma).

For quantification of EGFP fluorescence in IEC lineages, images were taken as 1um optical sections using a Zeiss LSM 700 confocal microscope. EGFP fluorescence intensity was measured in ImageJ. Briefly, CD326 was used as an epithelial-specific membrane marker and cells positive for lineage specific markers were outlined using the freehand drawing tool for pixel intensity measurement. Background signal was measured and subtracted from the raw value of the measured cells from the same image. Enteroendocrine cells were marked by CHGA (n = 36), tuft cells were marked by DCLK1 (n = 33), absorptive enterocytes were marked by FABP1 (n=97), Paneth cells were marked by LYZ (n = 78), goblet cells were marked by MUC2 (n = 62), intestinal stem cells (ISCs) were marked by OLFM4 (n = 78), and transit-amplifying progenitor cells were considered to be in the +4 position and not marked by OLFM4 (n = 78). Statistical significance was assessed by one-way ANOVA with post-hoc comparisons by Tukey’s test in Prism 8 (GraphPad).

For OLFM4+ and KI67+ cells in Id3^RFP/RFP^ and wild-type control mice, 1um optical sections were acquired using a Zeiss LSM 700 confocal microscope to ensure accurate quantification. OLFM4+ cells were quantified and localized based on crypt position, designating the first cell position at the base of each crypt as “0”. KI67 was quantified by counting all bisbenzimide positive nuclei in each crypt and determining the percent KI67+ cells. Statistical significance was assessed by unpaired t-test in Prism 8 (GraphPad).

## Acknowledgements

The authors would like to thank Drs. Mei-fang Dai and Yuan Zhuang of Duke University for providing Id3^RFP^ mice. Brian Golitz and Noah Sciaky for technical support with sequencing and deconvolution of sequencing files. Qiancheng You and Dr. Chuan He of the University of Chicago for technical input and training on Nano-hmC-Seal. Dr. Pablo Arial and the Microscopy Services Laboratory in the Department of Pathology and Laboratory Medicine at UNC Chapel Hill for assistance with confocal microscopy. Drs. Bailey Zwarycz and Joseph Burclaff and Othmane Jadi for helpful conversations and critical reading of the manuscript. Dr. Terry Magnuson and members of the Magnuson lab at UNC Chapel Hill for helpful conversations and feedback.

## Funding

This work was funded by the National Institutes of Health (K01 DK111709 and R03 DK122111 to ADG; R01 DK091427 to STM), the American Gastroenterological Association (Research Scholar Award to ADG), the Center for Gastrointestinal Biology and Disease (P30 DK34987 to ADG), the Burroughs Wellcome Foundation (Collaborative Research Travel Grant to ADG), and the Department of Defense (W81XWH-19-1-0423 to JRR). The Microscopy Services Laboratory, Department of Pathology and Laboratory Medicine, is supported in part by P30 CA016086 Cancer Center Core Support Grant to the UNC Lineberger Comprehensive Cancer Center.

## Data availability

RNA-seq, Omni-ATAC-Seq, and Nano-hmC-Seal data are available in the Gene Expression Omnibus under accession number: GSE131442.

---

**Supplemental Figure 1:**
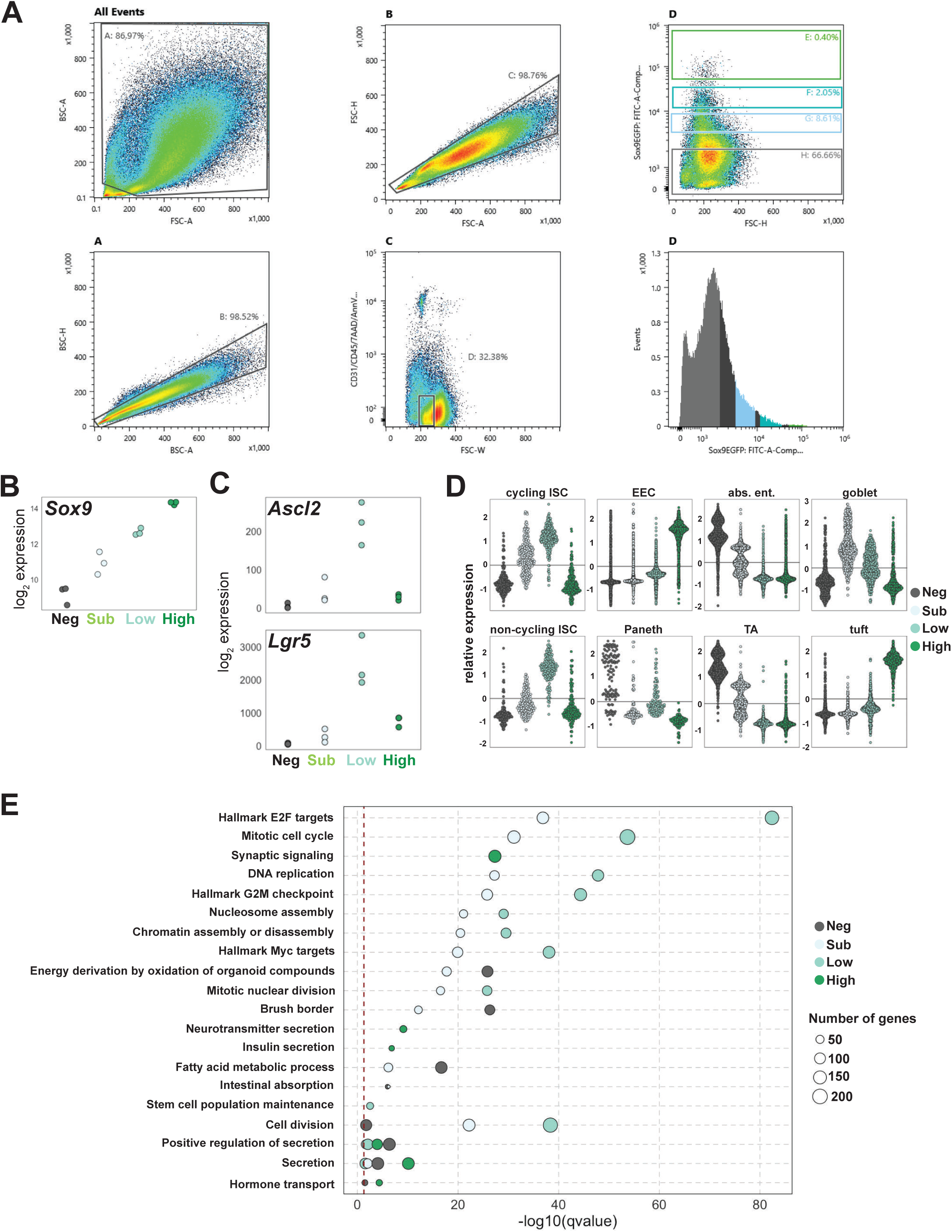
Transcriptomic characterization of Sox9^EGFP^ populations. (A) Representative gating for FACS isolation of Sox9^EGFP^ populations. (B) RNA-seq demonstrates accurate enrichment of endogenous *Sox9* mRNA across EGFP populations. (C) Stringent differential expression calling identifies some canonical ISC genes, such as *Ascl2*, which are enriched in *Sox9*^low^ cells only, and excludes *Lgr5*, which is upregulated in both *Sox9*^low^ and *Sox9*^high^ populations. (D) Bulk RNA-seq of Sox9^EGFP^ populations correlates with cell-type specific gene signatures identified by (Yan et al., 2017) (each point represents the expression of one gene). (E) GO analysis of Sox9^EGFP^ population transcriptomic signatures reveals enrichment of cellular functions consistent with predicted cell type. A subset of enriched categories are presented based on known relationships to IECs. The full analysis is presented in Supplemental Table 2.

**Supplemental Figure 2:**
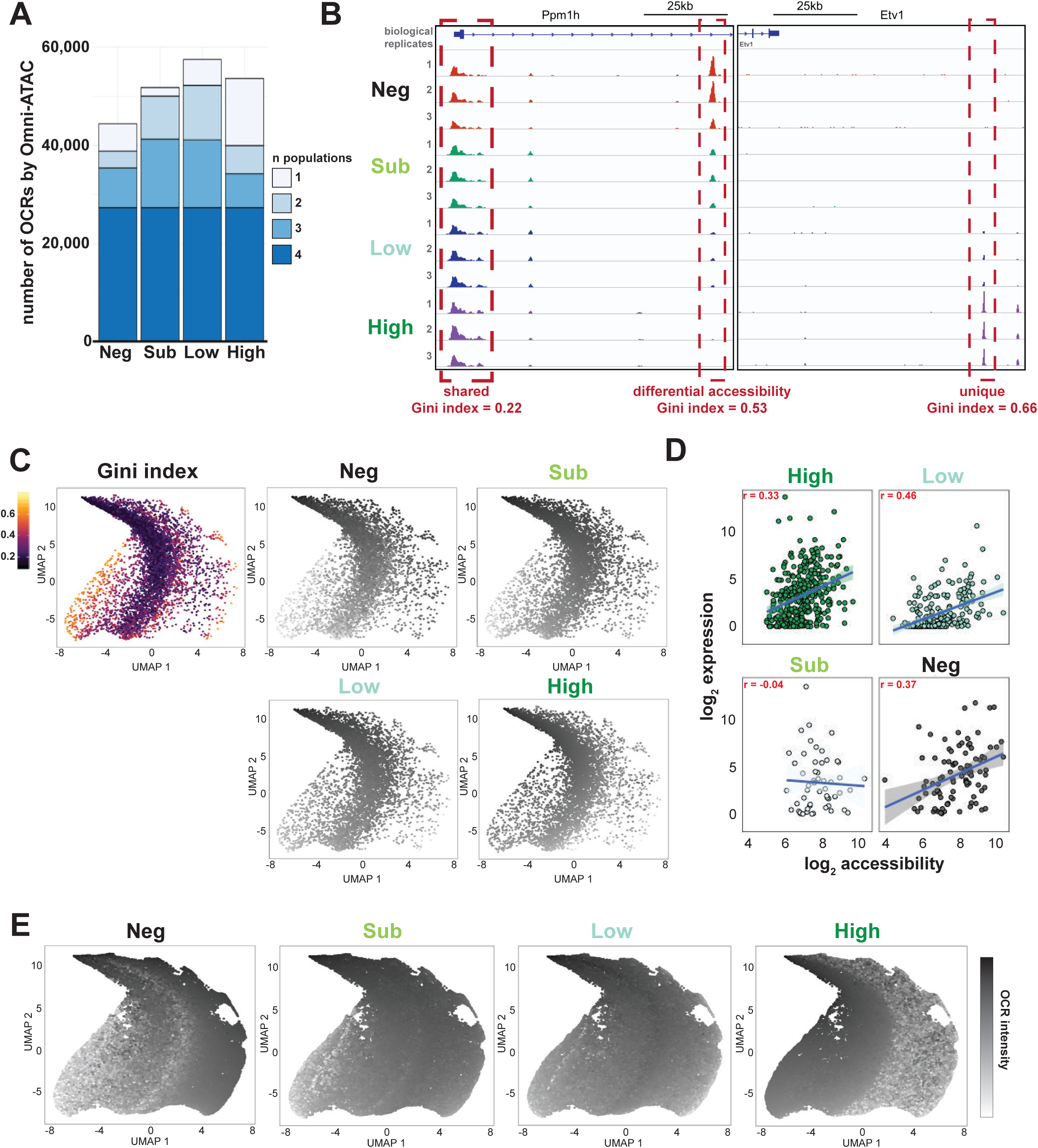
OCRs and promoter dynamics in IEC differentiation. (A) Summary of OCRs by Sox9^EGFP^ population demonstrates differential numbers of shared and unique peaks. Colors of bar graphs indicate the number of Sox9^EGFP^ populations in which a peak is observed. (B) Browser tracks show that Omni-ATAC-seq signal is highly reproducible between biological replicates and can be quantified by the Gini index to identify relative similarities and differences in OCRs between populations. (C) UMAP visualization of promoters reveals that a majority have low Gini index values indicative of high similarity between Sox9^EGFP^ populations, but also identifies dynamic promoter OCRs (n = 56 *Sox9*^neg^, 97 *Sox9*^sub^, 216 *Sox9*^low^, 341 *Sox9*^high^). (D) Correlation between accessibility and corresponding gene expression for dynamic promoters. (E) Sox9^EGFP^ population-specific OCR intensity reveals clustering by population on UMAP dimensionality reduction.

**Supplemental Figure 3:**
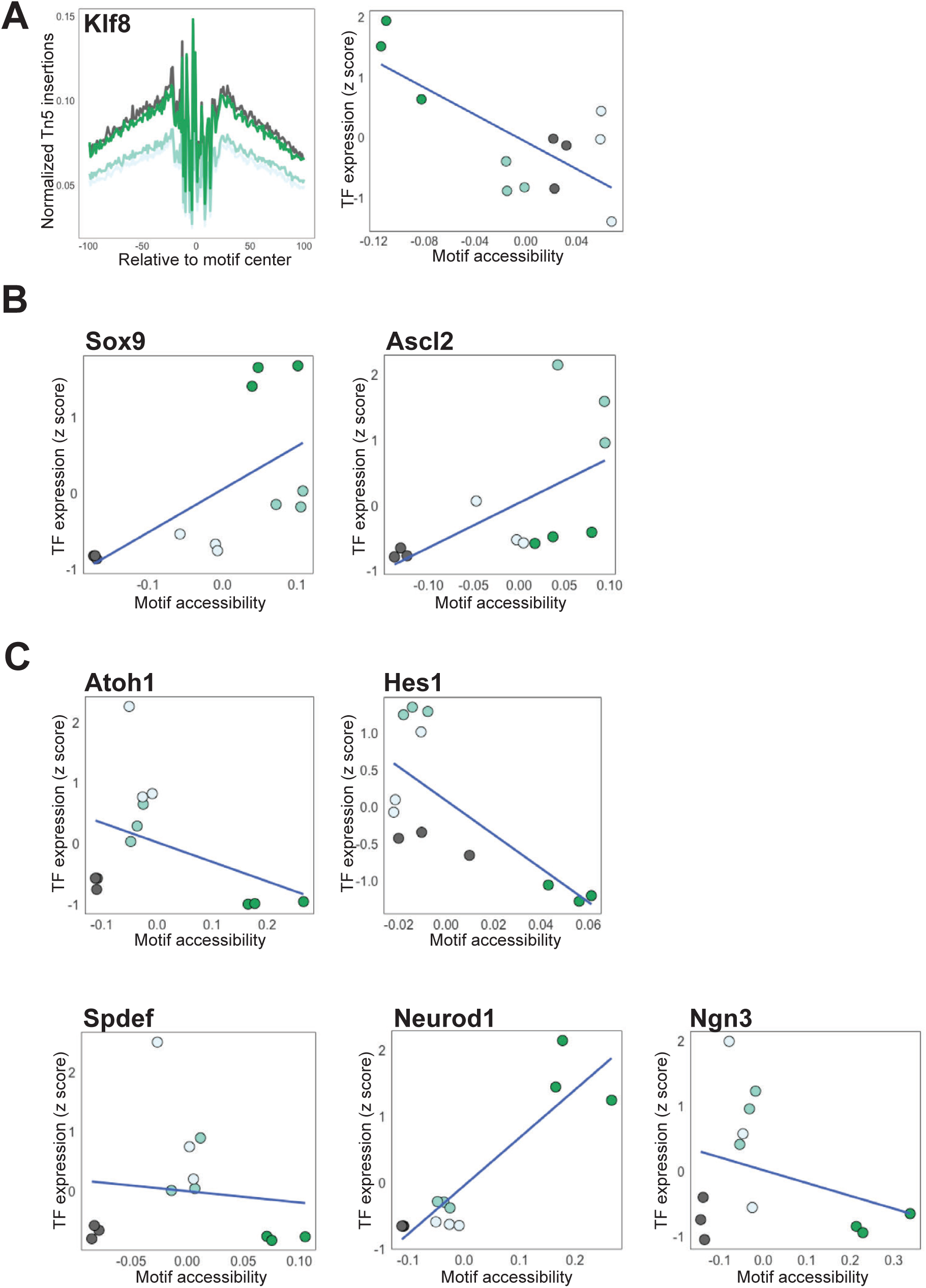
TF expression and motif accessibility identifies repressive relationships. (A) Tn5 insertion frequency relative to motif center (left) and TF expression plotted against motif accessibility (right) for *Klf8* demonstrates an inverse relationship between expression and motif accessibility predictive of *Sox9*^high^ specific repressive function. (B) Expression and motif accessibility relationships for ISC TFs *Sox9* and *Ascl2* demonstrate increased expression between *Sox9*^low^ and *Sox9*^high^ populations with little to no change in accessibility. (C) Relationships between expression and motif accessibility for canonical differentiation-associated TFs.

**Supplemental Figure 4:**
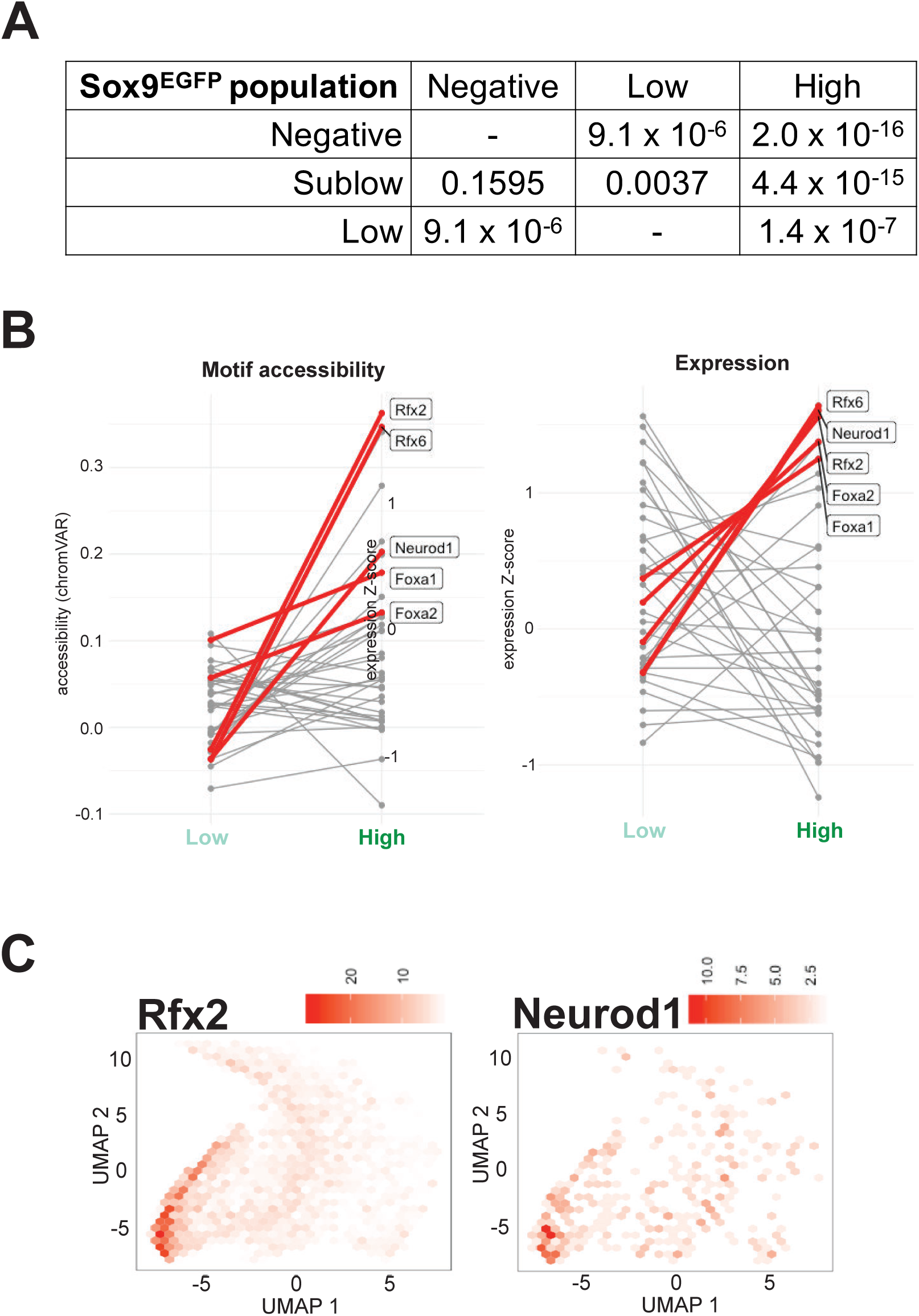
EEC TF expression and motif accessibility are dynamic in Sox9^low^ and Sox9^high^ shared OCRs. (A) Statistical analysis with pairwise comparisons for Fig. 5D, demonstrating significantly enriched expression of genes associated with *Sox9*^low/high^ shared OCRs in *Sox9*^low^ and *Sox9*^high^ populations. (B) Both motif accessibility and expression of EEC-associated TFs are increased in *Sox9*^high^ relative to *Sox9*^low^, consistent with differentiation. Plots show bias-normalized motif accessibility deviations from chromVAR (left) or z-scored RNA expression (right) in *Sox9*^low^ and *Sox9*^high^ populations and are identical to Fig. 5F with the exception of genes highlighted. (C) When mapped to UMAP plots, *Rfx2* and *Neurod1* motifs are distributed across *Sox9*^low^ and *Sox9*^high^ regions of the plot, but enriched in *Sox9*^high^ regions (n loci = 2,358 *Rfx2*; 959 *Neurod1*).

**Supplemental Figure 5:**
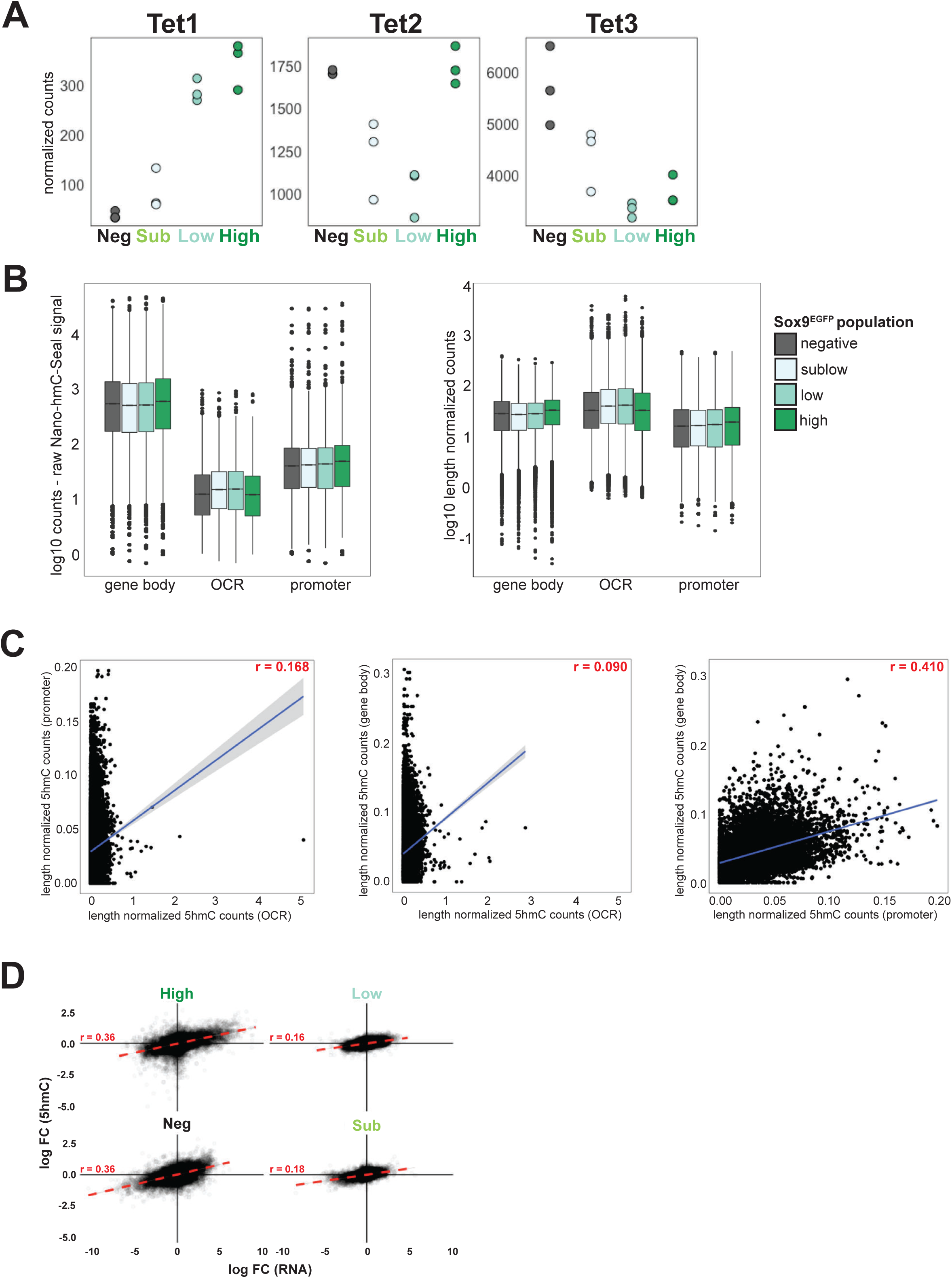
5-hydroxymethylcytosine distribution in IECs. (A) Normalized RNA-seq counts of *Tet1-3* show differential expression in Sox9^EGFP^ populations. (B) Raw Nano-hmC-Seal signal is enriched over gene bodies across all found Sox9^EGFP^ populations (left), but equivalent across gene bodies, OCRs, and promoters when normalized for feature length (right). (B) Correlation of 5hmC counts between different genomic classes show a positive relationship between promoter 5hmC levels and gene body hydroxymethylation (Pearson’s r is shown). (D) Relationship between log2 fold change of 5hmC and log2 fold change of expression for all genes (Pearson’s r is shown). Increased 5hmC in gene bodies demonstrates a moderate positive correlation with gene expression in all Sox9^EGFP^ populations, with stronger relative relationships found in *Sox9*^neg^ and *Sox9*^high^, consistent with increased DHMRs in these populations.

**Supplemental Figure 6:**
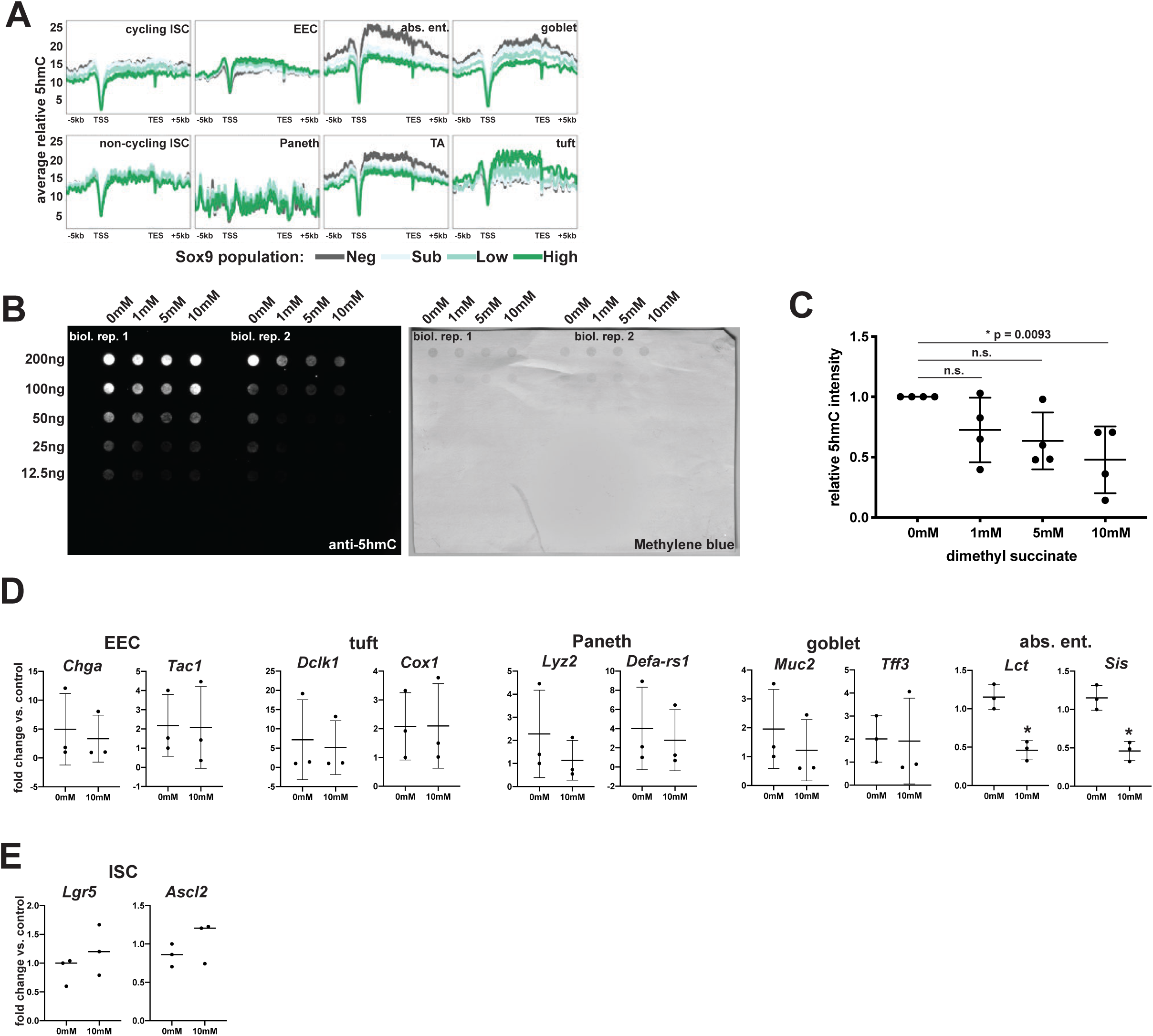
Modulation of 5-hydroxymethylcytosine in intestinal organoids. (A) 5hmC signal over scRNA-seq gene signatures from (Yan et al., 2017) independently confirms enrichment over absorptive enterocyte genes shown in Fig. 7A. (B) Representative 5hmC dot blot and loading control images from dimethyl succinate-treated intestinal organoids, two independent biological replicates. (C) Quantification of dot blot signal demonstrates significant decrease in global 5hmC with 10mM succinate treatment (n = 4 biological replicates per dose, samples normalized to their matched 0mM control, * indicates significance, p = 0.0093). (D) Five day treatment of organoids with dimethyl succinate demonstrates downregulation of absorptive enterocyte specific genes, with no change to canonical biomarkers associated with secretory lineages (n = 3 0mM and 10mM dimethyl succinate; * indicates p < 0.05, ** indicates p < 0.005). (E) Succinate treatment does not induce expression changes in ISC biomarkers *Ascl2* and *Lgr5* (n = 3 0mM and 10mM dimethyl succinate).

**Supplemental Table 1:** RNA-seq differential expression analysis

**Supplemental Table 2:** Significant GO terms for RNA-seq of Sox9^EGFP^ populations

**Supplemental Table 3:** Omni-ATAC peaks

**Supplemental Table 4:** chromVAR analysis of Omni-ATAC

**Supplemental Table 5:** Differentially hydroxymethylated regions

**Supplemental Table 6:** Primary antibodies

## Notes

The authors declare no conflicts of interest

